# Beta bursts carry a graded signal of semantic updating during naturalistic narrative comprehension

**DOI:** 10.64898/2026.09.27.754750

**Authors:** Abraham Goldstein, Yael Caspi

**Author notes:** **Corresponding author:** Abraham Goldstein.

## Abstract

Comprehending a narrative requires continuously updating a model of the discourse, and beta-band activity is thought to signal when that model should be held versus changed. Whether beta also grades how much to update, and whether that index is carried by discrete bursts beyond sustained power, is unknown for naturalistic comprehension. We recorded MEG from 61 human participants (22 men, 39 women) while they listened to naturalistic Hebrew narratives, detected beta bursts in the continuous signal, and quantified the semantic shift between successive sentences from sentence embeddings. Beta burst rate decreased after sentence boundaries, and regression-based deconvolution separated a pre-onset maintenance phase from a graded post-onset response whose magnitude scaled with semantic shift (peak ∼0.9 s). This response was independent of lexical surprisal, the acoustic envelope, and referent introduction, largely robust to dependency structure, though it shared variance with propositional quantity. Burst occurrence carried a component of the effect beyond the accompanying amplitude change: applying that change as a pure gain to each participant’s envelope and re-running the identical detection predicts only 75% of the observed burst-rate reduction (excess p = .038), the effect survived an amplitude-invariant threshold, and a Poisson model reproduced it as a graded reduction in burst-initiation rate. The effect held in both participant groups, which heard differently framed narratives, and was decisively supported (BF10 > 250). Beta bursts thus provide a graded signal of semantic updating during continuous comprehension, expressed in the rate of burst occurrence over and above the accompanying change in sustained power.

**Significance Statement:** Comprehending speech requires continuously updating an internal model of the unfolding discourse, but how the brain signals how much to update is unclear. Beta activity is thought to maintain the current cognitive state; we show that during natural story listening, the rate of brief beta bursts decreases at sentence transitions in proportion to how much the meaning shifts, consistent with a graded “how-much-to-update” signal rather than a binary boundary flag. Converging analyses show the effect is expressed in how often bursts occur, by more than the accompanying change in sustained beta power can explain, extending the transient-beta-burst framework to naturalistic language. The signal appears automatic and tied to linguistic structure rather than to subjective event segmentation.

## Introduction

Understanding a story as it unfolds requires more than recognizing each word: the listener must continuously maintain a model of the discourse and revise it as the narrative moves on. A central problem for the neurobiology of comprehension is therefore not only when this internal model should be updated, but by how much: small adjustments within a coherent topic, larger revisions when the topic shifts. How the brain encodes the graded magnitude of such updates during natural language is not known.

One influential proposal assigns this maintenance-and-update function to beta-band (≈13–30 Hz) activity. On the “status quo” account, beta is expressed while the current sensorimotor or cognitive state is held in place and is reduced, or released, when that state must change (Engel and Fries, 2010). A beta decrease, on this view, is not a lapse in processing but a positive signal that the prevailing model is being revised. Laminar and connectivity work offers a candidate mechanism: within predictive-coding accounts (Rao and Ballard, 1999), alpha/beta rhythms carry the top-down signals that inhibit feedforward processing, so that a release of beta permits the feedforward cascade associated with updating (Bastos et al., 2020).

This framework has a direct linguistic instantiation. Beta power builds up over the course of grammatically legal sentences, consistent with maintenance of an incrementally constructed sentence model, and returns toward baseline at syntactic violations and unexpected continuations, when the model must be revised (Bastiaansen et al., 2010; Lewis et al., 2016). In naturalistic spoken-narrative MEG, beta tracks syntactic dependency structure beyond low-level features, rises while dependencies remain open, and shows a biphasic profile around the opening of new dependencies (an increase before, a decrease after), while, notably, not being driven by lexical surprisal (Zioga et al., 2023). These results link beta to the maintenance and revision of a linguistic model.

Two questions remain open. First, these demonstrations are categorical: violation versus no-violation, open versus resolved dependencies. Whether beta carries a graded update signal scaling with how much the model must change has not been tested for naturalistic comprehension, where the update at each sentence transition is a continuous quantity: the semantic distance between successive discourse states. Second, beta is increasingly understood not as a sustained rhythm but as a train of transient events (Sherman et al., 2016; Feingold et al., 2015; Lundqvist et al., 2016), whose rate dominates averaged beta power and behavior (Shin et al., 2017). Prior language-beta findings are expressed in averaged power, leaving open whether the effect is, mechanistically, a change in how often discrete bursts occur.

Here we address both questions. We recorded MEG while participants listened to naturalistic Hebrew narratives, detected beta bursts in the continuous recording, and quantified the semantic shift between consecutive sentences as the cosine distance between sentence embeddings from a Hebrew neural language model (AlephBERT; Seker et al., 2022). Using regression-based deconvolution, we separated the graded semantic component of the burst-rate response from the generic word response, the binary boundary response, and the contributions of lexical surprisal and the acoustic envelope.

From the status-quo account we derived three confirmatory hypotheses: (H1) beta burst rate falls after sentence boundaries, consistent with release of a maintained model; (H2) the magnitude of this post-onset release scales parametrically with semantic shift, dissociably from lexical surprisal and the acoustic envelope; and (H3) the graded effect is expressed in burst occurrence over and above the accompanying amplitude change: the burst-rate modulation exceeds what that amplitude change mechanically produces through a threshold, and is robust to an amplitude-invariant threshold.

Beyond these confirmatory tests we include a planned specificity control (whether the graded signal is redundant with a boundary-locked P200/N400 evoked response) and two exploratory analyses: the cortical sources of the detection and gradient signals, and whether the linguistic boundaries coincide with subjective event boundaries (Zacks et al., 2007; Kurby and Zacks, 2008; Baldassano et al., 2017).

## Materials and Methods

### Participants

Sixty-one right-handed native Hebrew-speaking adults (aged 18–33 years, M = 24.25; 22 men, 39 women; 30 Approach, 31 Avoidance condition; five excluded from an initial 66 for missing/unusable data) provided informed consent under approval of the Ethics Committee, Department of Psychology, Bar-Ilan University (No. 04/18). These MEG recordings were acquired for, and have been reported in, a study of inter-subject neural synchrony during the same narratives (Caspi et al., 2026). That report concerns source-level alpha and beta synchrony between listeners; the beta-burst detection, boundary-locked deconvolution and all analyses reported here are new and do not overlap with it.

### Stimuli and procedure

Participants passively listened to two of four naturalistic Hebrew narratives (∼3 min each), written in the second person and recorded by a single male narrator, forming a 2 (emotional theme: anger, fear) × 2 (motivational framing: approach, avoidance) design. Because the same narrator recorded all four, speaker characteristics are constant across conditions. Emotional theme varied within participant (each heard one anger-themed and one fear-themed narrative, in counterbalanced order), whereas motivational framing was manipulated between participants (approach group n = 30, avoidance group n = 31). The two framing groups therefore heard non-overlapping narrative texts, although the underlying scenarios are partially crossed. An exam/cheating scenario appears in the fear-approach and anger-avoidance narratives and a deadline/technical-failure scenario in the anger-approach and fear-avoidance narratives, so each group heard one of each. Scenario content is therefore matched across the framing groups. In an independent sample of 70 raters the anger scripts elicited more anger than the fear scripts (t(69) = 8.79, p < .001) and the fear scripts more fear (t(69) = 2.46, p = .017), with no difference between the approach and avoidance framings on either rating. The two groups also did not differ in rated task motivation (t(58) = 0.22, p = .83; Caspi et al., 2026).

### MEG acquisition and preprocessing

MEG was recorded with a 4D/BTi 248-channel whole-head magnetometer system at 1017.25 Hz and 0.1-400 Hz band-pass filtering. Reference coils located above the sensors enabled removal of environmental noise. The head shape was digitized manually for subsequent source estimation. Five coils were attached to the head to record head positions and motion throughout the experiment. Line noise, building vibration and cardiac artifacts were removed following Tal and Abeles (2013), and a 10 ms audio-delay correction aligned the recordings to stimulus onset (Caspi et al., 2026). Analyses were restricted to magnetometers. Per recording, noisy channels (SD > 4000 fT) were detected and interpolated, and ocular/muscle artifacts were removed by ICA (FastICA, 25 components, fixed seed; Hyvärinen and Oja, 2000) fit on 1–40 Hz data; EOG-correlated (z > 2.0) or myogenic components were projected out. Cleaning was performed on the continuous recording; for boundary-locked epoch analyses, epochs were additionally cleaned with AutoReject (Jas et al., 2017; 5-fold CV; 6000 fT fallback). Preprocessing and sensor-level analyses used MNE-Python (Gramfort et al., 2013). Because ICA projection is a linear operation that removes only the EOG-correlated or myogenic components (typically a small subset of the 25) and leaves the remaining sources intact, the broadband morphology and amplitude of beta transients are preserved; all burst metrics were computed on the post-cleaning signal.

### Stimulus features

Narratives were transcribed and force-aligned with WhisperX (Bain et al., 2023). Sentence boundaries were defined as the first word of each transcript segment (sentence-level units from transcript punctuation); 208 of 210 segments (99%) ended in sentence-terminal punctuation (period or question mark). The WhisperX transcript was verified identical to the original story texts and the segment onsets force-aligned to the audio. Boundaries therefore reflect the authored sentence punctuation of the scripts rather than model-inserted segmentation, and the automatic step is the alignment of words to audio, not the placement of the boundaries. Authored punctuation is nonetheless a writing convention rather than a validated linguistic unit, and no independent hand-annotation was performed to confirm that these boundaries coincide with the units a linguist would mark. For each boundary, semantic shift was the cosine distance between AlephBERT sentence embeddings of consecutive segments (imvladikon/sentence-transformers-alephbert; Seker et al., 2022; Reimers and Gurevych, 2019), a continuous index of discourse-state change. Lexical surprisal (bits) was computed per word from a Hebrew GPT-Neo model (Norod78/hebrew-gpt_neo-small; Black et al., 2021); the broadband acoustic envelope (and pitch, spectral flux) was extracted from the speech. For the syntactic and lexical-semantic control battery, transcripts were parsed with Stanza 1.13 (Qi et al., 2020) using its Hebrew pipeline: tokenization, multi-word-token expansion (required because Hebrew fuses clitics orthographically), POS tagging, lemmatization and Universal-Dependencies dependency parsing, together with Hebrew named-entity recognition (IAHLT model), yielding per-boundary dependency length, tree depth, clause and verb counts, content-word turnover (new content lemmas relative to the previous sentence), and named-entity (referent) introduction; each was co-modeled as a competing parametric boundary regressor alongside semantic shift, and shared variance partitioned by commonality analysis.

### Beta-burst detection

Cleaned data were band-pass filtered 12–25 Hz and the magnitude of the Hilbert transform taken as the analytic amplitude envelope per channel. Bursts were periods exceeding the 70th percentile of each channel’s amplitude distribution for ≥100 ms; shorter crossings were discarded. This percentile-threshold-plus-minimum-duration criterion follows Tinkhauser et al. (2017), who likewise excluded events shorter than 100 ms, roughly two beta cycles, to limit the contribution of noise-driven amplitude fluctuations, and who swept a comparable range of percentile thresholds. The identical algorithm was applied to continuous data, event epochs, and source ROI time courses. Robustness of the burst definition was assessed by re-detecting bursts across eight amplitude thresholds (50th–90th percentile), five minimum durations (50–150 ms), and a wider detection band (13–30 Hz), and re-computing the boundary burst-rate decrease at each amplitude threshold, and the controlled semantic-shift response across the 60th to 80th percentile range, the five minimum durations and the wider band (N = 61; Supplementary Table S3); neither effect depended on these choices. Operationalizing beta as transient bursts and summarizing by burst rate follows the transient-beta-event framework: high-power beta arises as brief events, and the accumulation of these events across trials, i.e. event rate, is the dominant feature underlying behaviorally relevant differences in averaged beta power, so burst rate carries information partly obscured by trial-averaged power (Shin et al., 2017; for review see Lundqvist et al., 2024).

### Continuous burst-rate signal and deconvolution

Cortical bursting was summarized as the burst rate, or occupancy: the mean of the binary in-burst indicator, taken across channels at each moment for the continuous timecourse and across time within a channel for per-channel summaries. Rectification makes channel-averaging valid where raw-MEG averaging would cancel. Occupancy is jointly determined by how often bursts occur and how long they last, so burst-onset rate is reported separately wherever occurrence is at issue. The continuous burst-rate signal (full span, 100 Hz, stimulus-aligned via per-word MEG timestamps) was modeled by regression-based deconvolution (the regression-ERP or temporal-response-function framework, rERP/TRF; Smith and Kutas, 2015; Ehinger and Dimigen, 2019) to disentangle responses to events recurring every ∼260 ms. Each regressor’s response was expanded over lags −0.5 to +1.0 s with 10 raised-cosine basis functions. The estimated response to a given regressor, as a function of lag from the event, is referred to throughout as that regressor’s kernel: it is the time course the model attributes to that event type once the overlapping contributions of all the others have been accounted for. The design comprised word onset, word×surprisal, boundary onset, boundary×semantic-shift, and the continuous acoustic envelope; columns were standardized and fit per participant by ridge regression (penalty 1.0).

Effect sizes are Cohen’s d of the mean kernel amplitude in a 0.4–1.0 s window, a data-driven interval bracketing the post-onset release peak (∼0.9 s; see Results) while excluding the pre-onset maintenance elevation. To localize what about the sentence the graded release tracks, the shift modulator was recomputed from sentence openings of increasing length (first 3, first 5 words) and from noise-matched first/second sentence halves (∼4 tokens each), and the deconvolution refit with each variant. Because the shift is a distance between two sentences, it was additionally decomposed into quantities belonging to a single sentence (the incoming and the outgoing sentence’s cosine distance from the mean sentence embedding of that narrative) and into the shift values of the preceding and of the two following boundaries; each was co-modelled with, and separately substituted for, the canonical modulator.

### Burst-property decomposition and Poisson point-process model

To localize the shift effect to occurrence/amplitude/duration, the deconvolution was refit with the dependent signal replaced by burst-onset rate (initiations/bin) and by mean beta amplitude (channel-averaged envelope; conventional power). Event-level, within-participant correlations of semantic shift with post-onset (0.4–1.0 s) burst count, duration, and peak amplitude were tested across participants (Fisher-z). As a discrete-native confirmation, channel-pooled onset counts in 10 ms bins were fit per participant with a Poisson GLM (log link) on the same lagged basis. Binned-count Poisson regression of this form is the standard discrete-time approximation to a point-process model, in which covariates modulate the instantaneous rate of events, following the framework developed for neural spike trains (Truccolo et al., 2005); the shift kernel is therefore a log rate ratio per +1 SD shift. A peri-boundary onset histogram (PETH) split at the participant-wise median shift was computed from event times. Because burst rate is a thresholded function of the envelope, a reduction in amplitude lowers it mechanically; we therefore measured that mechanical relation rather than assuming it. Each participant’s per-channel beta envelope was multiplied by gains from 0.80 to 1.20 and the identical detection re-run with the threshold fixed at its unscaled value, giving the gain exponent (elasticity) dlog(rate)/dlog(gain) at unit gain, that is, the percentage change in burst rate produced by a one percent change in amplitude. The rate modulation predicted by a pure amplitude change is then that exponent times the observed fractional amplitude modulation, and the excess of the observed over the predicted modulation was tested across participants.

### Boundary-locked, time-frequency, source, and evoked analyses

Boundary epochs (−1 to +2 s) and 1:1 matched non-boundary control epochs (>3 s from any boundary) were contrasted on PRE (−0.8–0 s) vs late-POST (0.4–1.0 s) burst rate, and major-vs-minor used a within-subject median split on shift, major boundaries being those with above-median semantic shift and minor boundaries those below. Because high-shift boundaries had lower pre-onset (tonic) burst rates, a residualized analysis regressed late-POST on PRE burst rate within participant (pooling boundary and control epochs) to remove baseline differences, and the graded effect was re-tested on this residualized measure. Its dissociation from acoustic (pause duration, acoustic envelope, pitch), lexical (surprisal), motivational-framing (approach/avoidance), and sentence-length (segment word count) factors was assessed by within-participant multiple/partial regression and 2×2 ANOVAs (median splits). Pause durations were the forced-aligned inter-word intervals. Single-trial power (Morlet, 4–35 Hz) was summarized in theta/alpha/low-beta/high-beta sub-bands and a theta/beta ratio. Source projection used LCMV beamforming (Van Veen et al., 1997) with individualized single-shell BEM head models, built by warping a template MRI (MNI Colin27) to each participant’s digitized head shape (FieldTrip, Oostenveld et al., 2011; Caspi et al., 2026; ∼3273 sources/participant; 5% shrinkage; max-power orientation; unit-noise-gain weight normalization, which compensates for the depth-dependent bias in beamformer output; rank reduction enabled; matrix inversion), parcellated into 72 AAL ROIs (Tzourio-Mazoyer et al., 2002); per-ROI boundary-vs-control suppression and residualized burst×shift correlation (N = 61 for both) were corrected for multiple comparisons across ROIs by the false discovery rate (Benjamini and Hochberg, 1995). To test the laterality crossover as an interaction rather than as two separate contrasts, both effects were also computed from the same 61 participants’ source epochs, with detection indexed by the boundary PRE-to-POST change, for which no control epochs are needed. Each participant’s 72-ROI map was z-scored across ROIs before the hemisphere × effect interaction was tested across participants, the standard correction for comparing the topography of effects that differ in overall magnitude (McCarthy and Wood, 1985); this removes magnitude and retains spatial distribution, which is what a laterality claim concerns. A scale-free laterality index on the unscaled values and a permutation test shuffling hemisphere labels within homologous ROI pairs served as convergent checks.

Because burst rate is a per-ROI relative measure (proportion of time above each ROI’s own amplitude percentile) and all source effects are expressed as between-participant Cohen’s d, the ROI maps are insensitive to absolute differences in source amplitude or depth. For the evoked response, raw MEG was re-epoched (1–30 Hz, −1 to +2 s, baseline −0.2–0 s, top-50 channels) around boundaries and matched control words; P200 (200–300 ms) and N400 (300–500 ms; Kutas and Federmeier, 2011) were contrasted (paired t; cluster permutation, 5000 perms, |t| > 2.0) and correlated single-trial with shift and with late burst rate.

### Human segmentation (boundary-condition)

To test whether the sentence boundaries that drive the beta signal correspond to subjective event boundaries, an independent online study (JATOS) had Hebrew readers segment the written narratives into coarse-grained events by clicking the space between two words. Readers were asked to mark the points at which, in their judgement, one event unit ends and a new one begins, and were explicitly instructed to segment coarsely: to mark the largest units that seemed natural and meaningful, dividing each story into a relatively small number of large parts, each time a new central stage of the story began. They were told that a new event may begin when the action, place, time, characters or goal changes, that there are no rigid rules and no right or wrong answers, and they completed a short practice passage first. Raters read the narratives whereas the MEG participants heard them, so the human boundaries were established without prosodic cues. Valid raters (≥1 boundary) were retained; fine-grained outliers were excluded by a robust rule (per-reader median marks-per-story exceeding the group median by more than three scaled median-absolute-deviations), removing readers who segmented at fine grain (median 22–106 marks/story) against a group median of ∼6, i.e. who did not follow the coarse-grain instruction.

Consensus boundaries were locations endorsed by ≥50% of valid raters (marks within ±2 words pooled). Each AlephBERT boundary was assigned a graded human-agreement value (number of valid raters marking within ±2 words). We then (i) compared consensus boundaries to AlephBERT boundaries (recall/precision), (ii) tested whether human agreement related to semantic shift, (iii) re-locked the continuous burst-rate analysis to human-endorsed vs. human-rejected boundaries, and (iv) added human agreement as a parametric boundary regressor to the deconvolution (a parallel model; the canonical model was retained unchanged). N = 31 sessions (30 with ≥1 boundary; 27 valid after excluding one zero-mark reader and three fine-grained outliers), coarse grain only. The 27 valid raters were 19 women and 8 men aged 21–54 (M = 27.9, SD = 6.7); 25 reported Hebrew as a native language.

### Statistical inference

#### Experimental Design and Statistical Analysis

The experiment used a 2 (emotional theme: anger, fear; within-participant) × 2 (motivational framing: approach, avoidance; between-participants; approach group n = 30, avoidance group n = 31) design. Unless otherwise stated the participant was the unit of analysis (N = 61); analyses conducted at the level of individual boundaries (linear mixed models, variance partitions, single-trial evoked correlations) are identified as such and model participant as a random effect. For deconvolution and Poisson models, fit per participant, group inference used sign-flip cluster-based permutation (2000 perms, |t| > 2.0, cluster-mass; Maris and Oostenveld, 2007). Other contrasts used paired/one-sample t-tests on per-participant summaries; effect sizes are Cohen’s d. The occurrence-vs-amplitude comparison (H3) was tested directly: each dependent signal (burst-onset rate, occupancy, mean amplitude) was z-scored per recording before refitting the identical deconvolution, the 0.4–1.0 s shift-kernel window compared paired across participants (bootstrap CI and sign-flip permutation on the difference), the unique shift variance of each signal partitioned by commonality/semipartial analysis, and the rate effect re-estimated with the burst threshold fixed on non-boundary epochs (amplitude-invariant). To address non-independence of boundaries nested within narratives, the per-boundary residualized release was additionally modelled with a linear mixed model (participant random intercept and random slope for shift; narrative as a fixed factor), the shift×narrative interaction tested, and the shift kernel estimated separately for the two narratives as a two-item consistency check. Because the framing manipulation was between participants, each participant heard two of the four narratives, so narrative has two levels within a participant. A random effect estimated from two levels typically collapses to a singular fit, its variance being unidentifiable in practice, so narrative was treated as a fixed factor and per-narrative estimates are reported instead. Because motivational framing was a between-subjects manipulation, the shift × frame interaction was also tested and the deconvolution kernel estimated separately for the two framing groups as a between-subjects replication. Reliability was assessed by splitting boundaries into odd and even sets and testing the per-participant shift-release relationship in each half; Bayesian support was quantified with JZS Bayes factors (Rouder et al., 2009) for the shift-kernel amplitude and the Poisson log rate ratio, with a credible interval on the rate ratio.

### Data and Code Accessibility

All analysis code is publicly available at https://github.com/avigoldstein-biu/beta-bursts-semantic-updating, including the derived data and a script that recomputes every statistic reported here from those files; a permanent DOI will be minted on acceptance. The raw MEG data are available from the corresponding author (A.G.) upon reasonable request; the dataset is shared with Caspi et al. (2026), where it was first reported. The study was not preregistered.

## Results

### Beta burst rate decreases at sentence boundaries (H1)

Across 61 participants (206 unique sentence boundaries across the four narratives; 5,963 boundary and 5,859 control epochs), beta burst rate fell after sentence boundaries, from 9.6% to 8.6% (Fig. 1), an 11.0% reduction (boundary-vs-control change d = −0.70, t(60) = −5.42, p = 1.1×10⁻6), whereas matched control words showed no change (−0.05%). The time course showed this was not a sub-baseline dip but the decay of a pre-onset elevation: burst rate was elevated before boundary onset and declined thereafter, consistent with maintenance of the current model through sentence completion, followed by release. Control words were drawn from mid-sentence positions at least 3 s from any boundary, so they sample the trough of this cycle and sit below the sentence-final level throughout the epoch; the contrast is computed on the pre-to-post change in each condition, so this difference in level does not enter it. This biphasic profile, a pre-onset rise followed by a post-onset fall, matches the maintenance-then-update pattern reported for held-open syntactic dependencies in naturalistic MEG (Zioga et al., 2023) and motivates decomposing the response into components.

**Figure 1.**
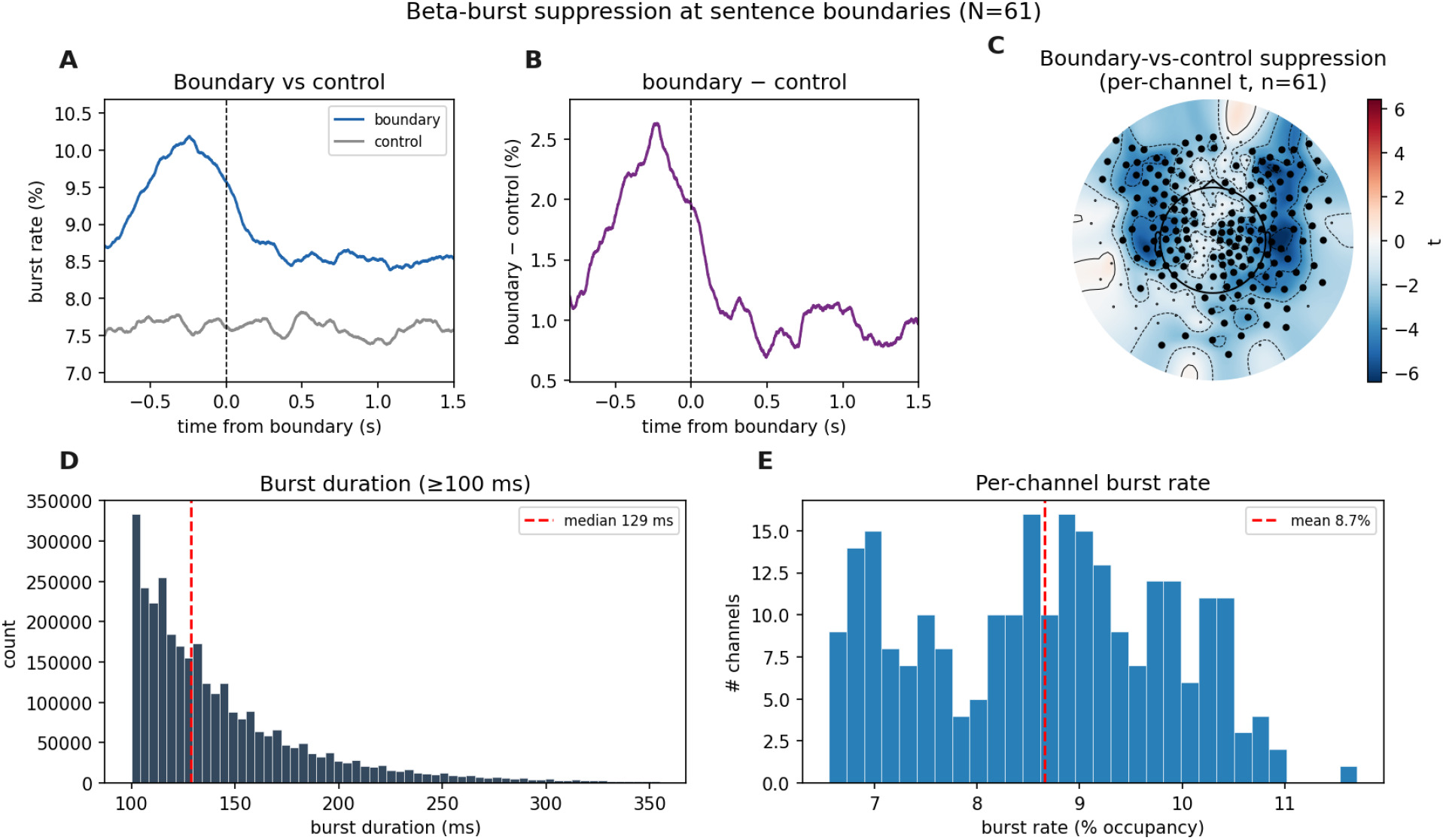
Beta-burst suppression at sentence boundaries (N=61). (A) Boundary vs. control burst-rate timecourse: a pre-onset elevation (10.2%) declining to ∼8.5% post-onset, against a control that is flat but lower throughout (mid-sentence words sample the trough of the same cycle); the contrast tests the pre-to-post change, not the level. (B) The boundary−control difference wave. (C) Suppression topography (per-channel boundary-vs-control t; 170/248 channels significant). (D) Burst duration under the analysis definition (12–25 Hz, per-clip 70th percentile, ≥100 ms; median 129 ms). (E) Per-channel burst rate (mean 8.7%, range 6.6–11.7%). The 70th-percentile amplitude criterion alone would place 30% of samples in a burst; requiring ≥100 ms of sustained supra-threshold amplitude reduces occupancy to ∼8.7%, so the spread across channels reflects variation in bursting rather than the position of the threshold.

### Deconvolution separates maintenance from a graded semantic release (H2)

Modeling the continuous burst rate as a sum of overlapping event responses separated the contributions of each regressor (Fig. 2); three are described here, with surprisal and the acoustic envelope below. The word kernel was a post-onset reduction (trough ∼0.4 s) with no evidence of an anticipatory rise (pre-onset d = −0.23, p = .08, a non-significant trend in the opposite direction to a pre-word increase). The boundary kernel carried a pronounced pre-onset elevation peaking ∼0.27 s before sentence onset (positive cluster, p < .001), a maintenance signal; the commonly reported “boundary suppression” is largely the decay of this elevation. The parametric semantic-shift kernel was a distinct late post-onset release peaking at +0.92 s (negative cluster, p < .001; 0.4–1.0 s d = −0.564), dissociable from both the generic word response and the boundary maintenance signal. The sign and the maintenance-to-release sequence are as predicted by the status-quo account (Engel and Fries, 2010; Lewis et al., 2016).

**Figure 2.**
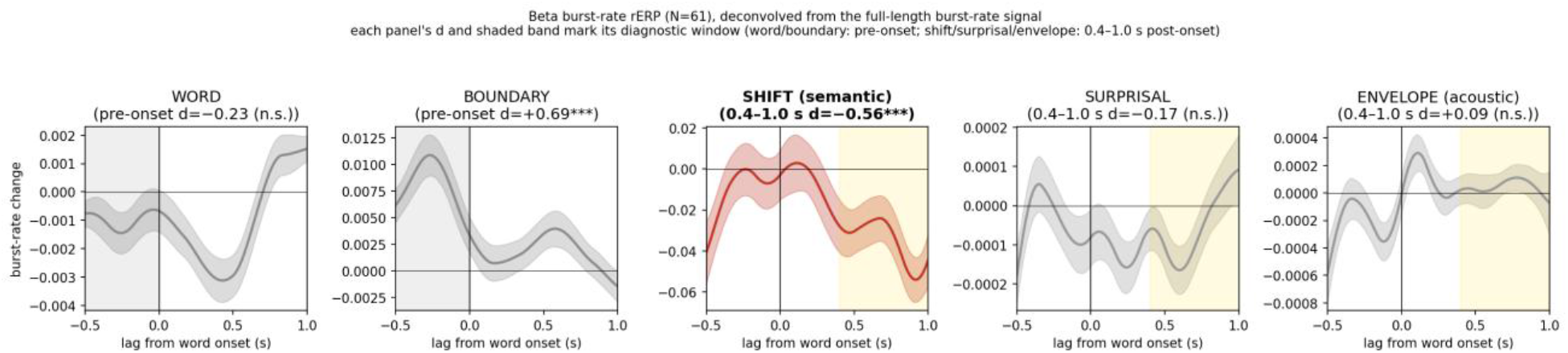
Deconvolution kernels (word, boundary, shift, surprisal, envelope). d values are Cohen’s d of the mean kernel amplitude in each kernel’s diagnostic window: the pre-onset window (−0.5–0 s) for the word and boundary kernels and the 0.4–1.0 s post-onset window for the shift, surprisal, and envelope kernels (shaded band in each panel). The word kernel shows no anticipatory pre-onset rise (d = −0.23, n.s.); the boundary kernel is dominated by a pre-onset maintenance elevation (d = +0.69, p < .001); the shift kernel is a graded post-onset release (d = −0.56, p < .001).

### The semantic release is graded, and is neither lexical nor acoustic (H2)

The shift kernel was estimated with lexical surprisal and the acoustic envelope co-modeled and survived unchanged (0.4–1.0 s d = −0.564, p < .0001) (Fig. 3), whereas the surprisal kernel was negligible and the envelope did not modulate the late window (d = +0.09, n.s.). It did not depend on how the burst-rate signal was constructed: refitting on a rate trace reconstructed from boundary epochs, rather than the continuous trace, reproduced the same kernel shape (r = 0.99). It converged across categorical (major-vs-minor d = −0.38), residualized-correlation (d = −0.45, p = .0009), and time-resolved (peak ∼0.95 s) analyses, all between d = −0.38 and −0.56 (Supplementary Figure S2). It was also invariant to the burst definition: across detection thresholds (60th–80th percentile), minimum durations (50–150 ms), and a wider band (13–30 Hz), the kernel held at d = −0.48 to −0.59 (peak ∼+0.9 s), cluster-significant in every case (Supplementary Table S3). An extended −1.0 to +2.0 s refit confirmed the peak was not edge-truncated (trough +0.93 s) and that the release was transient, significant over 0.7–1.1 s, recovering by ∼1.5 s. The signal therefore grades a discourse-timescale quantity, how much the topic changed, independent of lexical predictability or acoustic edges. The surprisal independence converges with Zioga et al. (2023), who found that lexical surprisal added nothing to beta beyond word frequency; we extend this from a categorical/syntactic to a graded semantic index (and confirm independence of word frequency itself; Supplementary Table S1).

**Figure 3.**
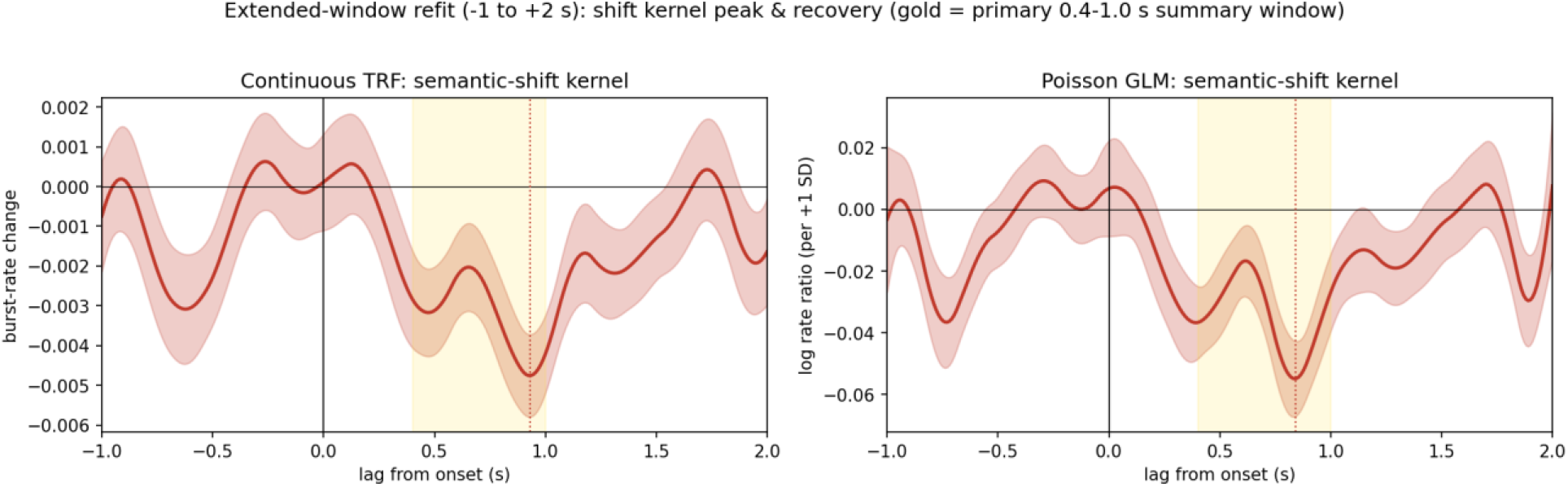
Extended-window (−1.0 to +2.0 s) refit of the semantic-shift kernel, from the continuous TRF (left) and the Poisson point-process model (right). The suppression peaks at ∼+0.93 s, is significant over 0.7–1.1 s, and recovers by ∼1.5 s, confirming that the peak is not truncated by the analysis window and that the release is transient. Shading is SEM; the gold band marks the primary 0.4–1.0 s summary window. N = 61.

### Residualized controls: the graded effect is not baseline, acoustic, lexical, or emotional

High-shift boundaries had lower pre-onset burst rates than low-shift boundaries, so a raw pre-to-post difference conflates tonic-baseline differences with boundary-evoked suppression. We therefore residualized late-POST burst rate on PRE rate within participant (pooling boundary and control epochs) and re-tested on this baseline-free measure. The graded effect survived cleanly: major boundaries showed greater residualized suppression than minor (d = −0.42, p = .002), residualized suppression scaled continuously with semantic shift (mean r = −0.041, d = −0.41, p = .002), and boundaries differed from matched controls (d = 0.53, p = .0001).

On this residualized measure, semantic shift dissociated from every competing predictor within a single framework. (i) Acoustics / pause. In a within-participant multiple regression (pause duration, semantic shift, envelope, pitch), semantic shift was the strongest predictor (d = −0.45, p = .0008), whereas pause duration (p = .67) and envelope (p = .39) did not contribute and pitch was only marginal (p = .049); a triple regression (shift + surprisal + pause) left shift as the sole predictor (d = −0.38, p = .004). The suppression thus occurs irrespective of whether the speaker paused, ruling out an acoustic-pause account. (ii) Lexical surprisal. Semantic shift predicted residualized suppression (d = −0.41, p = .002) but surprisal did not (d = −0.04, p = .79), the shift correlation being significantly stronger (t(60) = −2.24, p = .029); a 2×2 (shift × surprisal) showed a Shift main effect (F(1,60) = 7.45, p = .008) with no Surprisal effect (F = 0.74, p = .39) or interaction, and shift remained the sole predictor in a joint regression. (iii) Motivational framing. Motivation frame was a between-subjects manipulation; the graded effect did not differ between the approach and avoidance groups (shift × frame interaction p = .64; see Consistency across narratives and participant groups). (iv) Clause length. Semantic shift is confounded with sentence length (longer sentences carry lower shift; r = −0.43 within every participant); controlling for segment word count, the graded effect survived, partial-correlation d = −0.37 (from d = −0.41), p = .006, retaining ∼82–90% of its magnitude. In a combined per-epoch model the effect further survived covariates for narrative position, lexical frequency and word length, and a prosodic boundary-strength (pitch-reset) feature (all n.s.; shift d = −0.38, p = .004); only mean F0 level was marginally associated (p = .046), whereas the targeted pitch-reset measure was not, i.e. the marginal effect reflects overall pitch level, not boundary prosody (Supplementary Table S1). The graded post-boundary beta-burst suppression is therefore specific to discourse-level semantic change (Supplementary Figure S1): not a tonic-baseline artifact, an acoustic-pause or envelope response, a word-level prediction-error (surprisal) effect, or a function of the narratives’ emotional framing.

### The graded effect is dissociable from syntactic structure but shares variance with propositional quantity

Because the embedding cosine could co-vary with syntactic boundary strength, the quantity modelled by Zioga et al. (2023), we parsed the narratives with the Stanza Hebrew pipeline (Methods) and re-fit the deconvolution with each of a battery of interpretable boundary regressors added as a competing parametric modulator (Supplementary Table S2). The shift kernel was essentially unchanged by a dependency-length regressor, the held-open-dependency measure most analogous to Zioga (100% retained; the competitor carried no late kernel, d = −0.07, n.s.), and retained the majority of its magnitude against tree depth (77%), content-word turnover (81%), referent (named-entity) introduction (101%) and sentence length (78%). It was more substantially attenuated by clause count and verb count (to 52% and 37%), with which semantic shift is collinear (within-participant r ≈ −0.40 to −0.46; longer multi-clause segments carry smaller topic shifts). Co-modelling the full seven-regressor battery simultaneously attenuated the kernel further (42% retained, p = .067, not cluster-significant; Supplementary Table S2), the expected behavior when strongly collinear competitors are entered together.

Because co-modelled regression coefficients are unstable under this collinearity, we partitioned the variance of the per-boundary neural release directly: semantic shift retained a significant unique contribution after partialling clause count (semipartial p = .005) and verb count (p = .028), while clause and verb count contributed uniquely and in the opposite direction (more clauses, less suppression; p = .037 and .004). The graded release is therefore not reducible to syntactic clause/verb count, and is dissociable from dependency structure, but it is co-determined with the sentence’s propositional quantity, with which topic-shift magnitude is naturally correlated in these narratives.

### The release reflects an early topic property of the incoming sentence, not whole-sentence integration or completion of the previous one

We next asked whether the graded release tracks information the listener has actually heard by the time it occurs, or only the whole-sentence embedding, which necessarily includes words spoken after the response. By the time the release peaks (∼0.9 s), listeners have heard a median of four of the sentence’s ∼8 words. An embedding of only the sentence opening, available within the response window, predicted the release on its own: a first-five-word shift yielded a significant, graded kernel (d = −0.34, p = .010), and the effect declined monotonically as the opening was truncated further (first three words d = −0.14, n.s.). Because short-text embeddings are noisier indices of topic (their correlation with the full-sentence embedding falls from r ≈ 0.76 for five words to ≈ 0.55 for three), this attenuation tracks measurement reliability, not a loss of the underlying signal; the full-sentence embedding (d = −0.515 in this leading-edge model, the epoch-reconstructed counterpart of the −0.564 continuous fit) is therefore best read as the highest-SNR proxy for an early topic-divergence property, not as evidence that the response accumulates over the whole sentence. A noise-matched test confirmed this: splitting each sentence into first and second halves (∼4 tokens each), the two halves predicted the release equally (d = −0.391 vs −0.388), yet the second half is spoken after the 0.4–1.0 s window, so it can predict the release only through its correlation with the early-established topic. The graded release thus reflects a sentence-level topic-divergence property read early from sentence onset and best indexed by the whole-sentence embedding; it is neither temporal whole-sentence integration (the response does not wait for the sentence to finish) nor a preferential leading-edge effect.

Because a distance between two sentences is symmetric, however, a graded response to it is equally consistent with an update driven by the incoming sentence and with completion of the sentence that just ended. We therefore decomposed the shift into quantities belonging to a single sentence. The dissociation followed exactly which sentence each regressor contained. Regressors containing the incoming sentence produced the late release. Its atypicality for the narrative (cosine distance from the mean sentence embedding) predicted the release as strongly as the pairwise shift (d = −0.597, peak +0.86 s vs d = −0.564, peak +0.92 s). It also absorbed the pairwise term when both were modelled: shift retained 36% and was no longer cluster-significant, while the one-sided term carried its own late kernel (d = −0.380, p = .004).

Regressors that are properties of the preceding discourse alone (the outgoing sentence’s atypicality and the previous boundary’s shift, both fully determined before the boundary) do produce a late deflection when they stand in for the shift, as expected given their correlation with it (d = −0.39 and d = −0.28), but their kernels peak before boundary onset rather than in the release window, and once shift is co-modelled they carry no late kernel at all (d = −0.11, p = .38 and d = −0.15, p = .25) while the shift kernel remains intact (75% and 87% retained, cluster-significant). A shift regressor containing neither sentence (the boundary two positions later) was null (d = −0.135, p = .29, no cluster) and left the shift kernel at 100%, excluding leakage through the autocorrelation of the shift series. The graded release therefore belongs to the incoming sentence, while the preceding sentence modulates the pre-onset maintenance phase instead.

### The release is expressed specifically in burst occurrence (H3)

Five analyses localized the graded effect to burst occurrence and tested its relation to sustained power (Fig. 4). (i) Standardized contrast. Because burst-onset rate, occupancy and mean amplitude are measured on different scales, we z-scored each dependent signal before re-fitting the deconvolution, rendering the shift kernels comparable in within-signal SD units. In these units the occurrence kernel (onset rate, 0.4–1.0 s = 0.19 SD) and the amplitude kernel (0.26 SD) did not differ (paired Δ = −0.07, 95% CI [−0.15, +0.01], t(60) = −1.63, p = .11): burst occurrence is not modulated more strongly than amplitude. The larger raw Cohen’s d for onset rate (−0.59) than amplitude (−0.47) reflected its lower between-participant variance, not a larger effect. (ii) Unique variance. The two measures are, however, highly correlated (r ≈ 0.9), and partitioning their shared dependence on semantic shift revealed an asymmetry: the shift-dependence of occurrence remained significant after partialling amplitude (semipartial sr = −0.03, p = .013–.044 across raw and baseline-residualized variants), whereas the shift-dependence of amplitude did not survive partialling occurrence (p = .17–.72). The direct contrast between the two semipartials was not itself significant (paired t(60) = −1.69, p = .10), so we do not claim that occurrence carries more unique variance than amplitude, only that its contribution is the one reliably distinguishable from zero. The account in which the occurrence effect is a sustained-amplitude change viewed through a threshold is ruled out instead by (iii) and (iv). (iii) Excess over the amplitude prediction. Burst rate is a thresholded function of the envelope, so an amplitude reduction lowers it mechanically; the question is by how much. Scaling each participant’s per-channel envelope by a pure gain and re-running the identical detection gave a gain exponent of 4.12 ± 0.40 (occupancy) and 3.89 ± 0.39 (onset rate), the rate change produced by an amplitude change alone, duration criterion included. Multiplying each participant’s exponent by their observed fractional amplitude modulation predicts an occupancy modulation of −0.284, against an observed −0.405: the effect is 1.43× larger than a pure amplitude change produces (excess t(60) = −2.93, p = .005, d = −0.38). Burst-onset rate, the occurrence measure the claim concerns, showed the same excess at 1.34× (predicted −0.272 vs observed −0.363; t(60) = −2.12, p = .038, d = −0.27). The exponent is flat across gain ranges (4.07 at ±2% to 4.18 at ±20%), so this is insensitive to whether the amplitude change is spatially uniform or concentrated. Roughly 70% of the occupancy effect and 75% of the onset-rate effect is therefore what the amplitude change alone would produce; the remainder is not. (iv) Amplitude-invariant threshold. Re-detecting bursts with a per-channel criterion fixed on non-boundary (control) epochs, so the threshold cannot drift with the boundary-evoked amplitude change, left the effect intact (onset-rate d = −0.589, occupancy −0.570, peak +0.9 s, cluster p < .005); this addresses threshold drift specifically, not the mechanical dependence quantified in (iii). (v) Discrete model. A Poisson point-process model reproduced the effect with the appropriate noise model (shift kernel d = −0.543, peak +0.87 s): each +1 SD increase in shift reduced the burst-initiation rate to 0.965× (95% CI 0.950–0.982). Event-level analyses converged (shift predicted post-onset burst count d = −0.539, p = .0001, with weaker reductions in duration d = −0.292, p = .026, and peak amplitude d = −0.316, p = .016). These analyses show that the graded effect is expressed in the rate at which discrete bursts are initiated by more than the accompanying amplitude change can account for, even though amplitude is modulated to a comparable degree. The mechanistic claim is therefore a partial excess (occurrence carries a component beyond power), not magnitude (occurrence is not modulated more than power) and not independence (most of the rate effect is what the power change produces).

**Figure 4.**
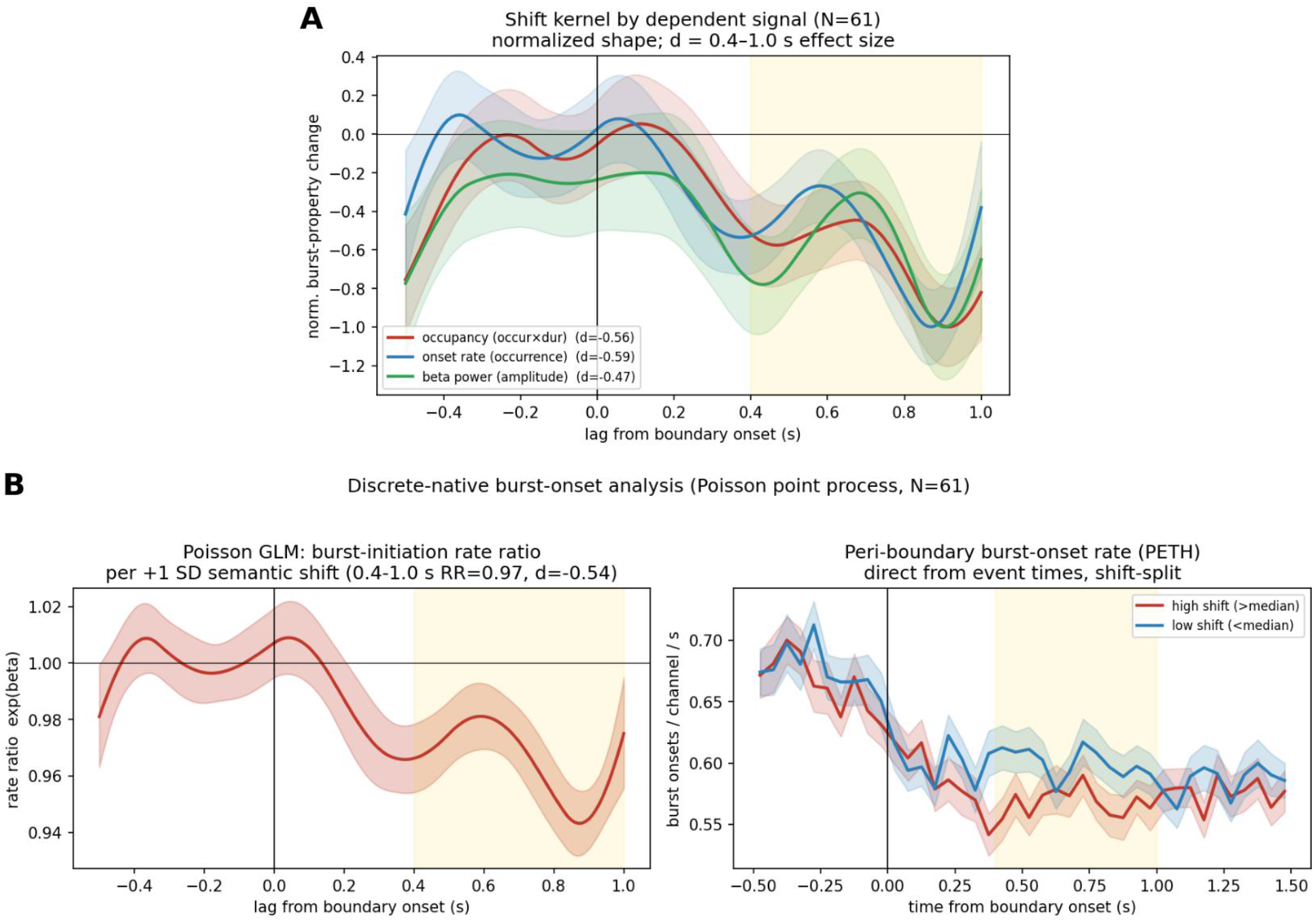
(A) Shift kernel by dependent signal, each curve normalized to its own peak: burst occurrence (onset rate), occupancy, and beta amplitude (power) share the same time course. Unstandardized effect sizes are d = −0.59, −0.56 and −0.47 respectively; these are not the basis of the occurrence-vs-amplitude comparison, because they differ mainly in between-participant variance. On the standardized within-signal scale used for that test the occurrence and amplitude kernels do not differ (Results). (B) Discrete Poisson point-process rate-ratio kernel (RR=0.97 per +1 SD shift) and peri-boundary onset histogram (PETH; high vs low shift).

To test whether the suppression selects specific burst shapes, individual beta-burst waveforms (662,411 bursts; 61 participants) were submitted to PCA. Shape variance was low-dimensional (8 PCs, 97%) but dominated by oscillatory phase (near-equal-variance PC1/PC2, PC3/PC4 pairs), a continuum rather than discrete burst subtypes, consistent with Rayson et al. (2026). Splitting bursts into waveform-type tertiles (PC1), all types showed significant post-boundary suppression (all p < 10⁻⁶) with no difference between them (ANOVA interaction F(2,120) = 0.97, p = .38), and shape-metric distributions were indistinguishable for boundary versus control bursts. The semantic suppression therefore operates uniformly across burst shapes, unlike the waveform-selective rate modulation reported for sensorimotor beta (Rayson et al., 2026), pointing to a general reduction in burst generation rather than a shape-selective gate, and reinforcing the burst-occurrence interpretation.

### Reliability of the graded effect

The graded effect reproduced in independent halves of the boundaries: split into odd and even boundaries, the per-participant shift-release relationship was significant in both (odd r = −0.050, p = .007; even r = −0.040, p = .033). This is the per-boundary estimator; re-fitting the full deconvolution within each half is noisier and reached significance in the odd half only (odd d = −0.62, p < .0001; even d = −0.16, p = .22). Bayesian evidence for the group effect was decisive, shift-kernel amplitude BF10 = 462, and the Poisson burst-initiation rate ratio RR = 0.965 (BF10 = 270) with a 95% credible interval [0.950, 0.982] excluding 1.0. Between-participant differences in kernel magnitude were not reliable (split-half r ≈ −0.09), consistent with a small but consistent group-level effect. Low between-participant variance is both what makes a within-participant effect robust at the group level and what limits the reliability of per-participant estimates (Hedge et al., 2018), so we make a group-level claim and do not interpret per-participant effect sizes.

### Consistency across narratives and participant groups

In a linear mixed model with a by-participant random slope, the graded effect held (shift β = −3.05, p = .0004) and survived adding narrative as a fixed factor (p = .0009); the shift×narrative interaction was null (p = .93), and the shift kernel estimated separately for the two narratives did not differ (anger d = −0.40, fear d = −0.31; paired p = .83). We treat this as two-item *consistency*, not item-level replication.

Because motivational framing was manipulated between participants, the two groups (n = 30 and n = 31) heard non-overlapping narrative pairs, permitting a stronger between-sample test. The graded release was present in both groups (approach β = −3.53, p = .013; avoidance β = −2.71, p = .011) with no shift × frame interaction (p = .64; p = .63 controlling emotional theme), and the deconvolution kernel replicated across groups (approach d = −0.58, p = .003; avoidance d = −0.54, p = .005; between-group difference p = .84). The effect is therefore consistent across two independent participant samples under different motivational framings, in addition to the two emotional themes.

At the single-narrative level all four narratives showed a same-signed (negative) effect, but with one ∼3-min narrative per cell only one of the four reached significance individually (per-boundary p: AngerApproach .065, AngerAvoidance .042, FearApproach .134, FearAvoidance .242; single-clip kernel d = −0.33, −0.51, −0.37, −0.26, respectively). Per-narrative cells are therefore individually underpowered, and the evidence for consistency rests on the between-groups comparison above rather than on any single narrative. The two groups’ kernels also differ in morphology: the avoidance-group kernel’s largest deflection fell pre-onset (≈ −0.5 s, driven by the anger-avoidance narrative) rather than at the canonical +0.9 s post-onset peak. What replicates across groups is therefore the windowed 0.4–1.0 s suppression, which is individually significant in each group and does not differ between them (p = .84), rather than the full shape of the kernel.

### The effect is specific to the beta band

Time-frequency analysis localized the suppression to beta (Fig. 5): the beta band (13–30 Hz) showed significant post-onset suppression peaking at ∼18–19 Hz (p < .0001; ≈ −4 to −7% depending on the post-onset window); alpha showed weaker partial suppression and theta a slight increase. The theta/beta ratio rose +8.9% at boundaries (p = .0015). In this unresidualized epoch-level comparison the ratio also tracked continuous shift (d = −0.35, p = .009) whereas the raw pre-to-post change in burst rate did not (d = +0.04, p = .77), which is the tonic-baseline confound described above and the reason the primary analyses residualize; on the residualized and deconvolved measures the burst-rate effect is d = −0.45 to −0.56. The effect is therefore predominantly beta rather than a broadband change, with theta moving in the opposite direction.

**Figure 5.**
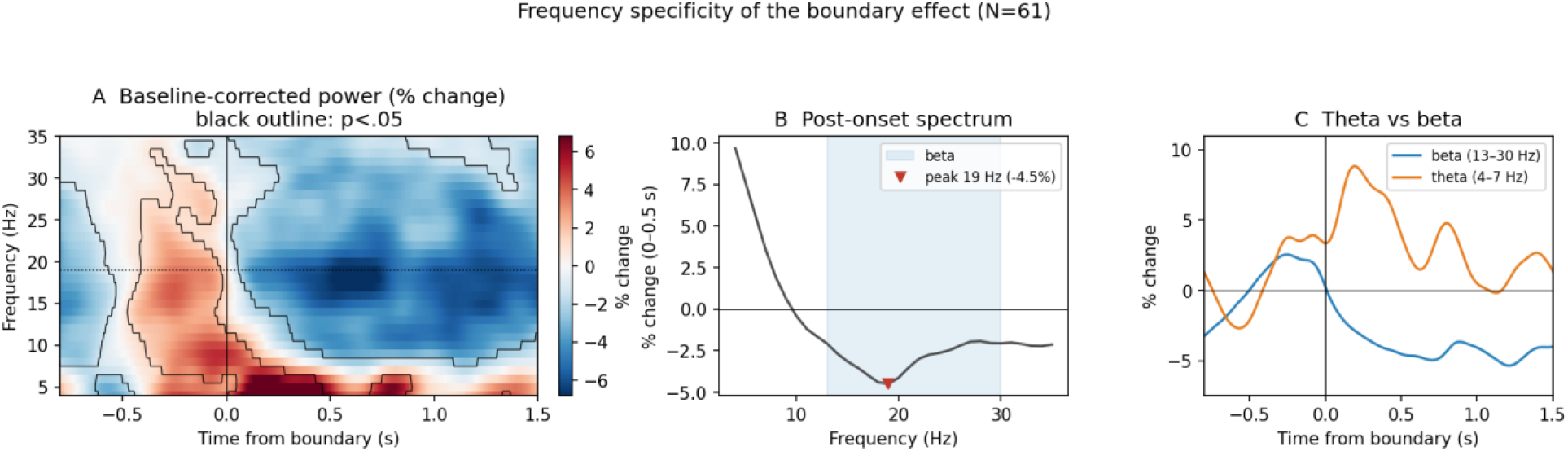
Frequency specificity (N=61). (A) Baseline-corrected time-frequency map (% change) with p<.05 outline, post-onset beta decrease, theta increase. (B) Post-onset (0–0.5 s) spectrum; beta trough at ∼18–19 Hz. (C) Opposing theta (increase) vs beta (decrease) time courses.

### Exploratory: a source-laterality dissociation between detection and grading

Projecting bursts to source space (unit-noise-gain LCMV with individualized single-shell BEM head models, 72 AAL ROIs) revealed a hemispheric crossover between detecting a boundary and grading it (Fig. 6). Boundary suppression (N = 61) was widespread and left-dominant (mean Left d = −0.90 vs Right −0.50): 67/72 ROIs FDR-significant (0/72 for controls), including left inferior/middle frontal cortex and Broca’s area (Frontal_Inf_Tri_L d = −1.16, Frontal_Mid_L −1.14, Insula_L −1.02) alongside posterior cortex. The semantic gradient (residualized burst×shift, N = 61) showed the opposite asymmetry, right-lateralized (mean Right d = −0.35 vs Left −0.19): FDR-significant in 24/36 right ROIs versus 0/36 left (24/72 total), with the major-vs-minor split converging (7 right vs 0 left). Because these were separate analyses on different samples, we tested the crossover directly, computing both effects from the same 61 participants’ epochs. The detection map reproduced the left-dominance (L −0.018 vs R −0.008, t(60) = −7.69, p = 1.6×10⁻10) and the gradient the right-dominance (t(60) = +2.72, p = .008). The hemisphere × effect interaction was reliable (t(60) = −10.24, p = 8.6×10⁻15, d = −1.31), as were a scale-free laterality index computed from the unscaled values (detection −0.43 vs gradient +0.17, t(60) = −7.17, p = 1.3×10⁻9) and a permutation test shuffling hemisphere labels within homologous ROI pairs (p < .0001). The two lateralities are opposite in sign rather than different in degree. The interpretable result is therefore the crossover, left hemisphere for that a boundary occurred, right hemisphere for how much the discourse changed.

**Figure 6.**
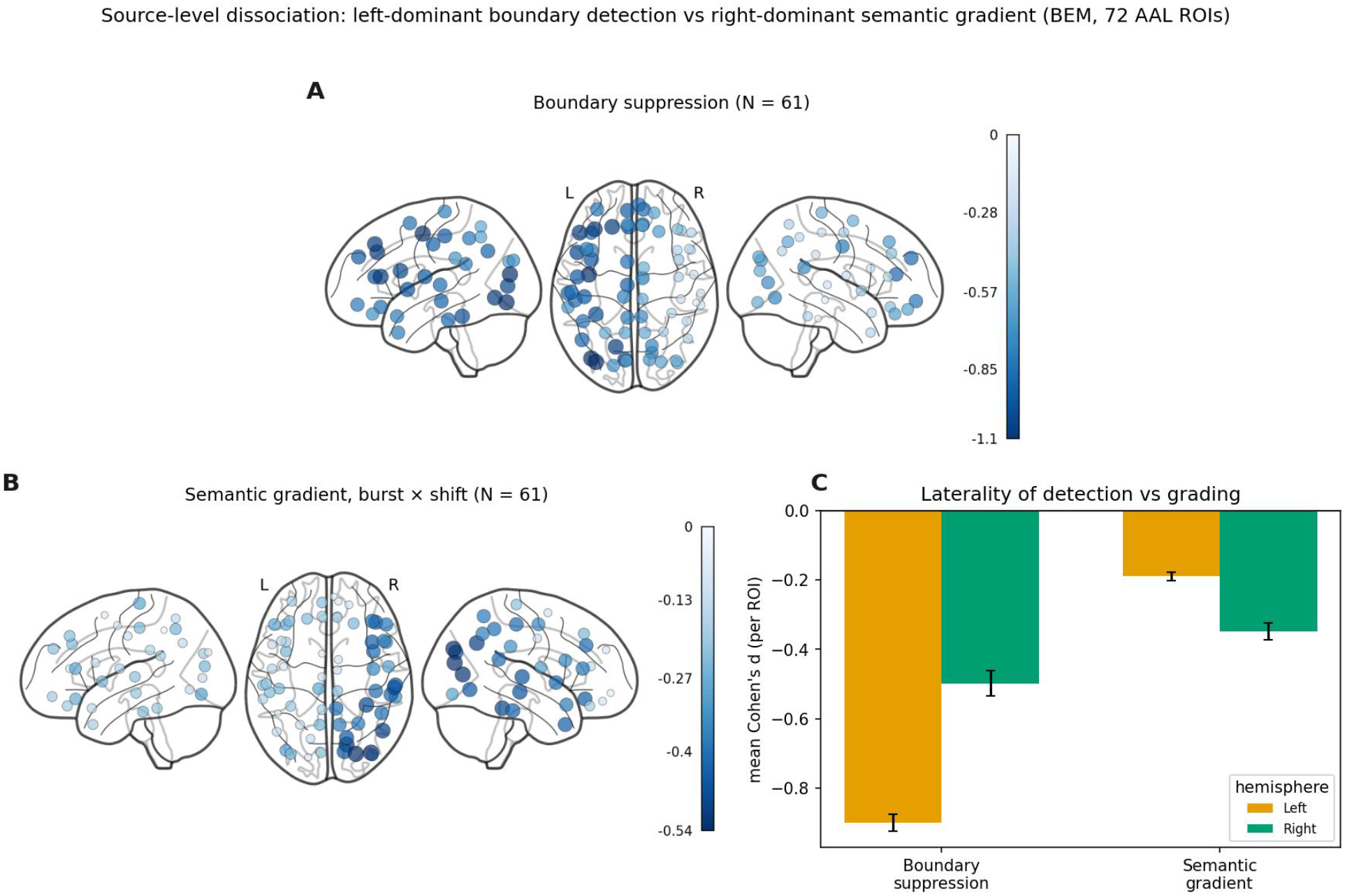
Source-level detection/gradient dissociation. (A) Boundary-suppression Cohen’s d per ROI (N=61), left-dominant (mean Left d=−0.90 vs Right −0.50). (B) Semantic-gradient (burst×shift) d per ROI (N=61), right-dominant (mean Right d=−0.35 vs Left −0.19). (C) Mean d by hemisphere for each effect: the laterality crossover.

### Specificity: the graded beta signal is not redundant with a boundary-locked evoked response

To test whether burst suppression merely re-describes an ERP (Fig. 7), boundary vs. matched control words were contrasted in the raw evoked response. N400 showed no difference (d = +0.12, p = .34); a more positive P200 appeared at boundaries (d = +0.33, p = .011) but only at trend level under cluster correction (243–298 ms, p = .084). Single-trial correlations dissociated the signals. Late beta burst rate uniquely tracked shift (r = −0.038, t(60) = −3.22, p = .002, d = −0.41), whereas no evoked component did (P200×shift r = +0.001, p = .94; N400×shift r = +0.016, p = .21), and the P200 did not differ between major and minor boundaries (d = +0.03, p = .83). Single-trial P200 amplitude also failed to predict the subsequent beta burst rate (P200–burst_post r = +0.026, p = .075; P200–burst_late r = +0.020, p = .16). The graded beta signal is therefore an independent channel rather than a downstream consequence of the evoked response. The evoked response marks boundary occurrence in a binary fashion; beta-burst release alone grades the magnitude of semantic change.

**Figure 7.**
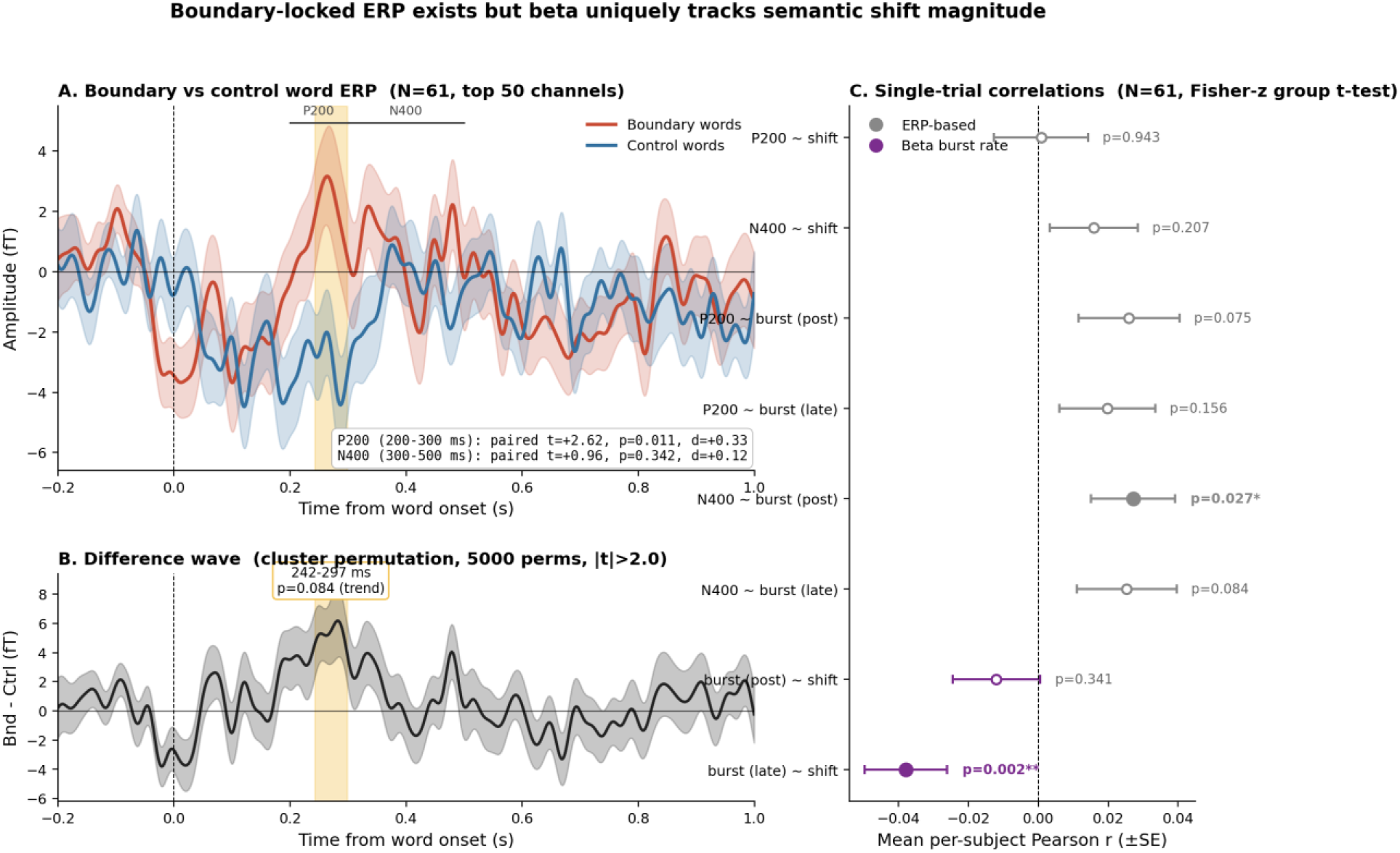
The graded beta signal is not redundant with the boundary-locked evoked response (N = 61, top 50 channels). (A) Grand-average ERP at sentence boundaries and matched control words; shading is SEM, bands mark the P200 (200–300 ms) and N400 (300–500 ms) windows. (B) Boundary−control difference wave with cluster permutation (5000 permutations, |t| > 2.0). (C) Single-trial correlations (per-participant Pearson r, Fisher-z transformed, group t-test): each ERP window against semantic shift and against burst rate, and burst rate against shift. Only late burst rate tracks semantic shift; no evoked measure does.

### Exploratory boundary-condition: these are linguistic, not subjective-event, boundaries

A separate human segmentation study established that the beta-driving boundaries are linguistic units rather than subjective event boundaries. Readers (27 valid) segmented each ∼480-word narrative into only a handful of coarse events (group-consensus boundaries: 1/3/4/3 across the four texts, 11 total), versus 44–56 AlephBERT sentence boundaries per text (206 total). Human consensus boundaries were a near-perfect subset of the AlephBERT boundaries (100% recall) but selected only ∼5% of them (precision ≈ 0.05), and the boundaries readers converged on were discourse/causal pivots (e.g. clause-initial “suddenly”, “because”) rather than the largest semantic shifts: human agreement was uncorrelated with semantic shift (Spearman ρ = −0.02, p = .75).

Two analyses show the beta signal is not contingent on subjective event status (Fig. 8). First, re-locking the continuous burst-rate analysis to human-rejected AlephBERT boundaries still produced robust suppression (dz ≈ −0.63), statistically indistinguishable from human-endorsed boundaries across a 0.33–0.67 consensus-threshold sweep (the endorsed-vs-rejected contrast was non-significant and unstable: dz = −0.12 to +0.18, all p ≥ .17). Second, adding graded human agreement as a parametric boundary regressor to the deconvolution (a parallel model; the canonical model retained) produced no reliable agreement kernel (0.4–1.0 s d = +0.02, p = .88), with moderate Bayesian support for the null (BF01 = 7.1), while the shift kernel was unchanged (d = −0.57). The null held across seven nested specifications (with/without shift, acoustics, word/surprisal; agreement d = +0.02 to +0.11, all p > .41; BF01 = 5.1–7.0): the graded agreement term adds nothing beyond the binary boundary and the semantic-shift terms. This null is specific to the tested post-onset window: in the same models the agreement kernel showed a pre-onset excursion (−0.4 to −0.1 s, d = −0.36, p = .006 in the canonical model, p < .05 in six of the seven specifications), in a window where the shift kernel was itself null (d = −0.06, p = .62). Because that window was identified after inspecting the kernels, we report this dissociation as an observation rather than a test. It is not, however, prosodic: adding per-boundary pause duration and pitch reset as competing modulators left the pre-onset agreement effect unchanged (d = −0.37, p = .006, 101% retained), although pause itself carried a pre-onset kernel of opposite sign (d = +0.31, p = .019). Whatever relates human agreement to beta therefore resides in the pre-onset maintenance phase, which the graded analysis removes as baseline, and its basis remains untested. At the level of raw per-boundary suppression, higher human agreement was weakly associated with stronger beta suppression (Spearman ρ = +0.20, p = .005; two-level agreement effect dz = −0.27, p = .04), but this does not survive as an independent post-onset kernel, even with semantic shift omitted from the model, consistent with human agreement marking cleaner sentence-boundary events rather than a graded event-salience signal carried by beta.

**Figure 8.**
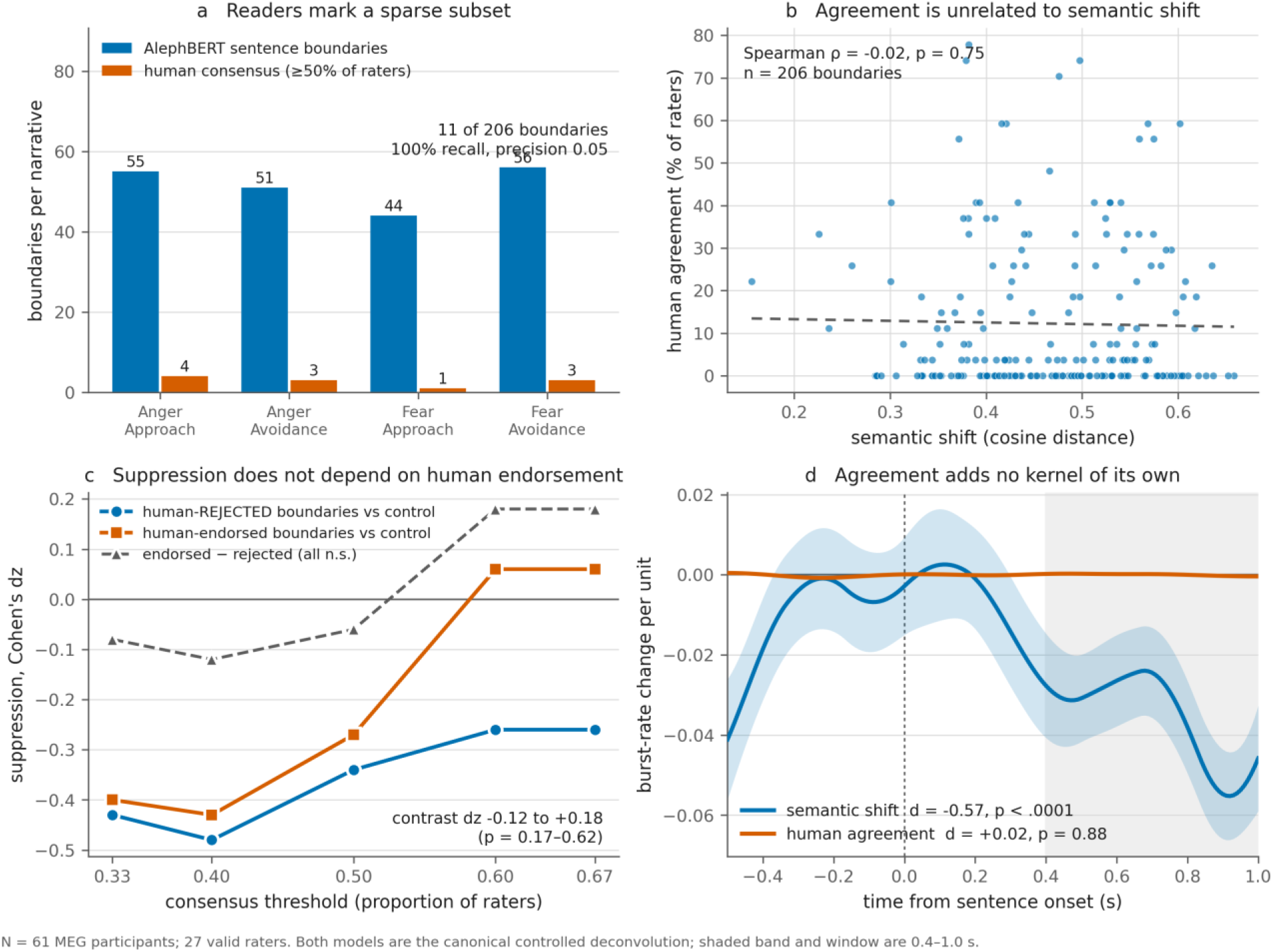
Human consensus vs. AlephBERT boundaries; agreement⊥shift; consensus-threshold sweep; agreement-augmented deconvolution kernels.

These results indicate that the beta release reflects an automatic linguistic-semantic update keyed to sentence structure and graded by semantic change, not a correlate of subjective event segmentation, which is sparser, discourse-cued, and uncorrelated with semantic-shift magnitude; graded human agreement contributes no independent component to the beta response beyond the sentence-boundary event itself.

## Discussion

Listening to a story, the brain must decide not only when its model of the discourse should change but how much. Beta bursts provide this graded signal. Burst rate decreased after sentence boundaries, and deconvolution isolated a graded post-onset release scaling with the semantic shift between successive sentences (peak ∼0.9 s, d ≈ −0.56), independent of lexical surprisal and the acoustic envelope. The effect was carried in part by how often bursts occur, the rate change exceeding what the amplitude change mechanically produces. Two exploratory analyses separated detection from grading, and the driving boundaries are linguistic, not subjective event boundaries.

### Extending the beta status-quo account

The sign and time course match the proposal that beta signals maintenance of the current cognitive state and is released when that state must change (Engel and Fries, 2010). In language, beta builds over grammatical sentences, drops at violations (Bastiaansen et al., 2010; Lewis et al., 2016), and tracks syntactic dependency structure (Zioga et al., 2023). We reproduce the pre-onset elevation, post-onset release and surprisal-independence, and extend them: where prior demonstrations are categorical and syntactic, ours is a continuous release graded by semantic change. The framework predicts that a change of state suppresses beta, not that suppression scales with its magnitude. That graded function is our first contribution, locating the signal at discourse-model updating rather than discrete syntactic events.

A syntactic regressor did not absorb the semantic kernel: the release was unchanged by dependency length and dissociable from tree depth, distinguishing it from the structure Zioga et al. (2023) modelled. It was partly shared with propositional quantity (clause and verb count), because high-shift boundaries open short, topically divergent sentences and low-shift boundaries longer elaborations. That coupling is a property of natural discourse rather than an artifact of our stimuli; the question is therefore whether each carries unique variance, and our partition shows both do. A deflationary account would attribute the release to processing demand, but the direction of the effect argues against it: sentences with more clauses and verbs showed less suppression rather than more (unique contributions opposite in sign to shift, p = .037 and .004), as did longer sentences and those with deeper trees, whereas demand accounts predict that more material produces a larger response.

### The topic shift is registered early

The release peaks ∼0.9 s after sentence onset, when only about the first half of the sentence has been heard, yet scales with a whole-sentence measure. This is not a contradiction: the previous sentence is complete, so only the direction of change need be estimated, and narrative sentences front-load their topic, so an embedding of the opening alone predicts the release. The release therefore reflects early registration of a topic shift rather than computation of a complete sentence representation, and re-locking to the second and third words shifted the trough progressively earlier rather than re-peaking per word, as a single onset-locked event predicts.

Decomposing the shift into one-sided quantities places the release on the incoming sentence: a measure of that sentence alone predicted the release as strongly as the pairwise distance and absorbed it, whereas measures of the preceding sentence carry no late kernel once shift is co-modelled. Because the pairwise distance is symmetric, this separates an update triggered by the new sentence from a wrap-up.

### Expressed in burst occurrence

The second contribution is mechanistic. Prior language-beta findings, including Zioga et al. (2023), are expressed in averaged power. Treating beta as a train of transient events (Shin et al., 2017), we asked whether the semantic effect is separable from sustained power. It is, partly: the observed rate modulation is ∼1.4× larger than applying that amplitude modulation as a pure gain and re-running the identical detection produces, it survives an amplitude-invariant threshold, and a Poisson model reproduces it with a discrete noise model. Because event rate rather than amplitude dominates averaged power, a language-beta effect may at base be a rate change. We frame this as a partial excess, not predominance: the residual ∼30% is what the burst framing adds.

A beta decrease with state updating fits the status-quo account and predictive-routing accounts in which alpha/beta inhibit feedforward signaling, so releasing beta permits the update cascade (Bastos et al., 2020). The direction is not uniform: Sedley et al. (2016) report the opposite sign, beta power correlating positively with the magnitude of perceptual prediction updates, though in a different measure and domain. Beta also rises while a structure is held open (Zioga et al., 2023), which matches our pre-onset elevation, and with content reactivation (Spitzer and Haegens, 2017), so the release is compatible with a reactivation reading (Rose et al., 2016) as well as a status-quo one. The effect is dissociated from lexical surprisal, the canonical proxy for prediction error, though beta can track surprisal in continuous speech (Weissbart et al., 2020), so this is specificity rather than insensitivity: a large topic shift need not be a large prediction error.

### Detecting a boundary vs. grading its magnitude

Two exploratory analyses suggest the brain separately registers that a boundary occurred and how much changed. Boundary suppression was left-dominant whereas the semantic gradient was right-lateralized. The crossover is a tested interaction rather than two contrasts falling on opposite sides (hemisphere × effect, d = −1.31). We interpret it only hemispherically, since the strongest gradient ROIs fall in occipito-parietal cortex where localization is least reliable. It is not a reflection of the dataset’s general beta topography: in these same recordings beta synchrony between listeners is left-dominant (Caspi et al., 2026), so the right-dominant gradient runs counter to the prevailing laterality, consistent with right-hemisphere discourse integration (Jung-Beeman, 2005; Youssofzadeh et al., 2022). In the evoked response, a boundary-locked positivity marked occurrence but did not scale with shift, whereas single-trial late burst rate uniquely tracked it.

### Linguistic, not subjective event boundaries

A natural reading would invoke event-segmentation theory, in which comprehenders segment at points of prediction failure and neural event structure is recoverable from narrative (Zacks et al., 2007; Kurby and Zacks, 2008; Baldassano et al., 2017). An independent segmentation study argues against that reading here. Readers marked far fewer events than sentence boundaries, at discourse and causal pivots rather than the largest semantic shifts, and agreement was uncorrelated with shift; suppression was indistinguishable at endorsed and ignored boundaries, and adding graded human agreement produced no reliable post-onset kernel while the shift kernel was unchanged, with moderate Bayesian support for that null (BF01 = 5.1–7.1). The release therefore appears to reflect an automatic linguistic-semantic update keyed to sentence structure rather than subjective event segmentation. The two are nested rather than rival: the boundaries readers converge on were a subset of the sentence boundaries (100% recall, ∼5% precision), so a sparse, consciously accessible event structure sits within a dense, automatic one, and we locate the beta signal at a continuously updated discourse model.

These recordings were first reported as a study of inter-subject synchrony (Caspi et al., 2026). There, approach motivation raised beta synchrony between listeners; here, the graded release is unchanged by motivational framing (shift × frame p = .64), so motivation modulates the shared, sustained beta regime rather than the boundary-locked response. That study would not be expected to show this response: inter-subject correlation (Hasson et al., 2004) indexes envelope similarity in 5 s windows and cannot resolve a 0.9 s peak.

### Limitations

Several caveats bound these conclusions. (i) Our mechanistic claim is non-redundancy, not magnitude: amplitude is modulated comparably to occurrence (paired Δ = −0.07, n.s.), so we claim only that the rate change exceeds what the amplitude change mechanically produces, by ∼35–45% (onset rate p = .038, occupancy p = .005). The effect is small (≈ 3.5% rate change; residualized r ≈ −0.04); it is decisively supported (BF10 > 250) and replicates across data halves at the group level, but per-participant magnitudes are not reliable. (ii) Stimuli were four ∼3-min Hebrew narratives (2 × 2 theme × framing) under passive listening; we modelled the non-independence explicitly and found a between-groups replication with consistent sign across all four, but per-narrative cells are underpowered and generalization across languages, genres, and tasks is untested. (iii) Semantic shift is an embedding-based distance, not a validated measure of discourse-model updating or prediction error, and correlates with propositional quantity, which cannot be fully decorrelated within four narratives. (iv) The human boundary condition rests on 27 readers at a single, coarse grain, so it bounds the relation to coarse event boundaries rather than to human segmentation generally, and boundaries were defined automatically from authored punctuation. (v) The sign tension with Sedley et al. (2016) remains unresolved. (vi) The source localization is exploratory and hemispheric-level only (single-shell BEM from a template MRI warped to each head shape, unit-noise-gain beamformer); spatial leakage precludes claims about specific generators. (vii) All effects are defined at sentence transitions; semantic change within a sentence, and discourse in which sentence boundaries are not well defined, lie outside what we tested.

Beta bursts thus extend the status-quo account from a categorical to a parametric update law.

## Author contributions

Y.C. and A.G. designed research; Y.C. performed research (data curation); A.G. analyzed data; A.G. wrote the paper; A.G. and Y.C. edited the paper.

## Acknowledgments

The authors used Claude Code (Anthropic) to assist in writing the analysis and figure-generation code, and in manuscript preparation and language editing. All analyses were specified, reviewed, and verified by the authors, who take full responsibility for the content of this article.

---

**Figure S1.**
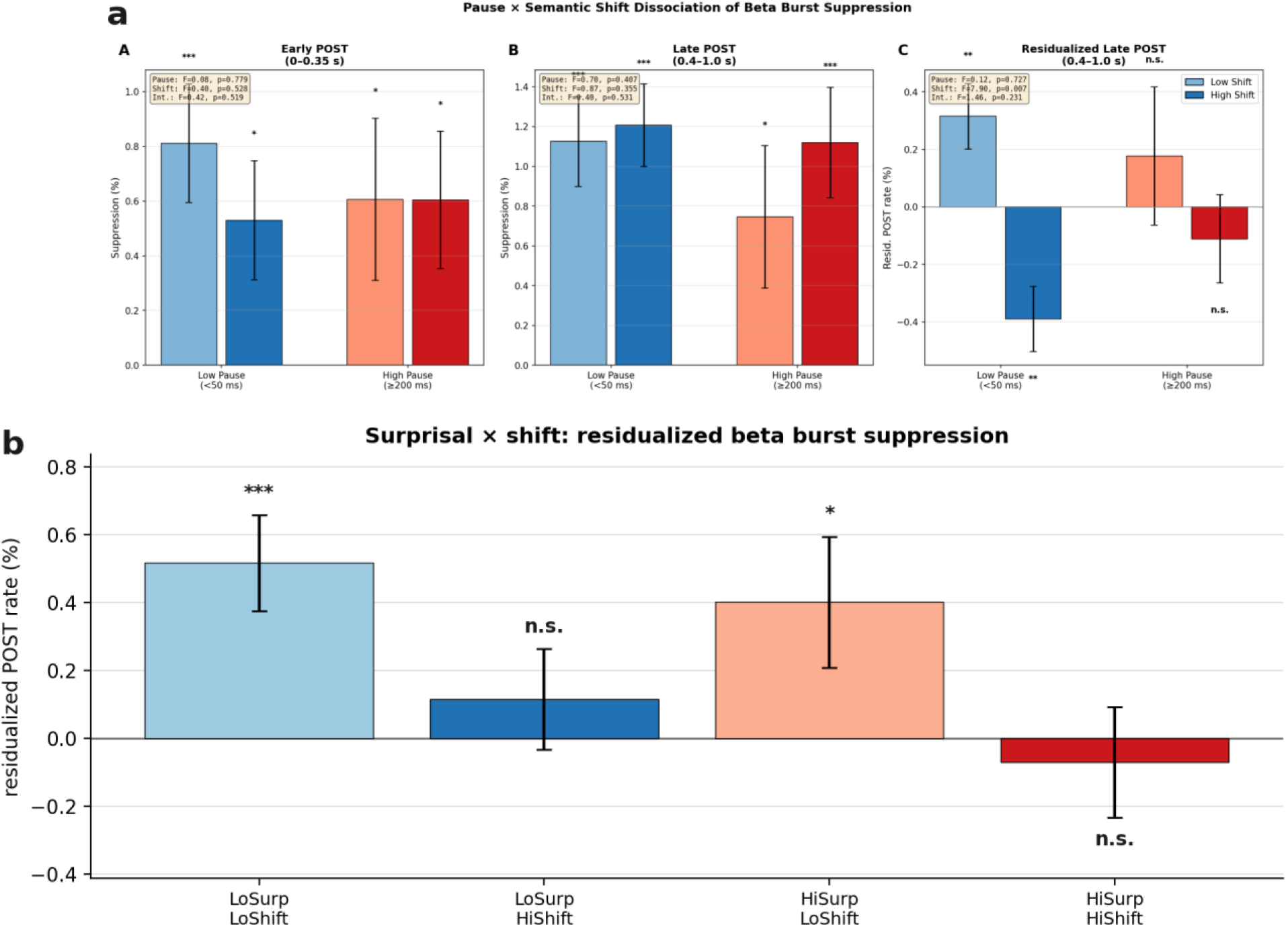
Dissociations of beta-burst suppression. (A) Pause × semantic-shift, for the early post-onset window (0–0.35 s), the late window (0.4–1.0 s), and the late window residualized on the pre-onset baseline. (B) Surprisal × semantic-shift on the residualized late window.Median splits; N = 61; error bars are SEM.

**Figure S2.**
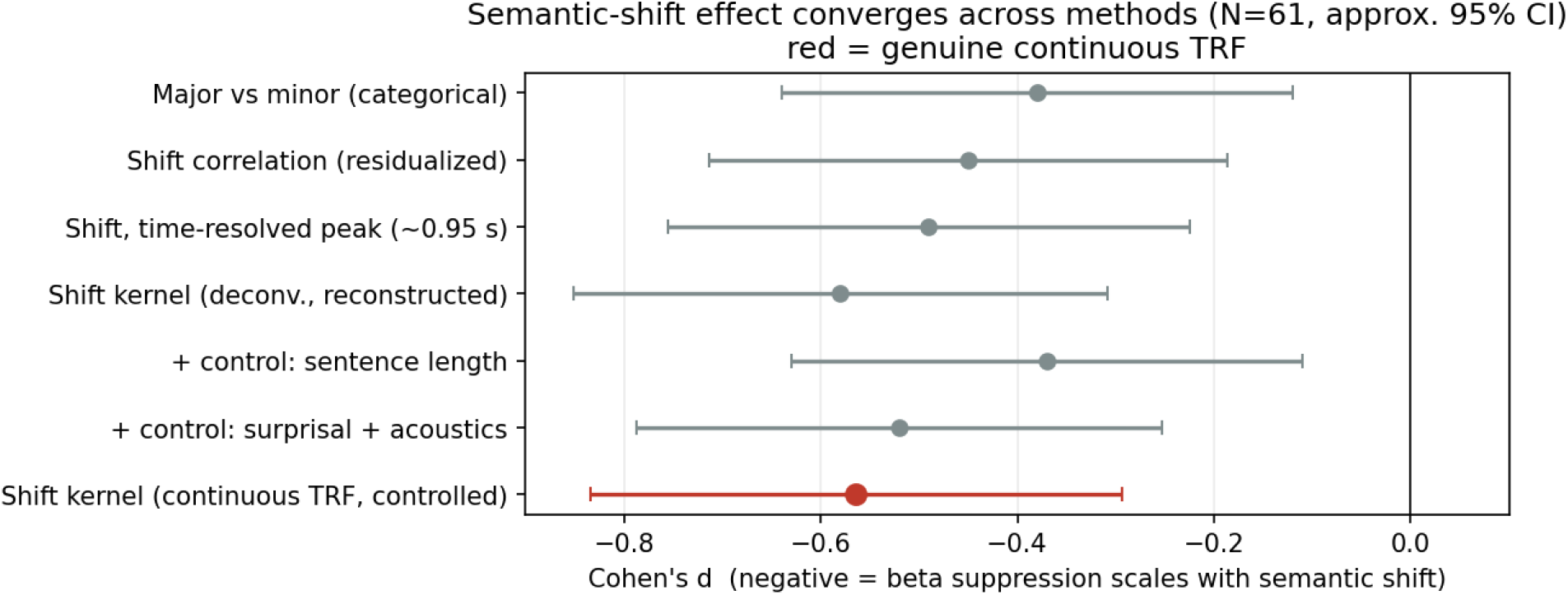
The semantic-shift effect converges across analysis methods (N = 61, approximate 95% CIs). Cohen’s d in the 0.4–1.0 s window for the categorical major-vs-minor contrast, the residualized shift correlation, the time-resolved peak, the epoch-reconstructed kernel, two co-modelled controls, and the canonical continuous-TRF kernel (red). Negative values indicate that beta suppression scales with semantic shift.

**Supplementary Table S1.** Combined covariate control on residualized suppression. Per-epoch residualized late-POST burst rate regressed on semantic shift and eight covariates (standardized β, per-participant OLS, group one-sample t-test; N = 61 participants, 5,880 boundary epochs). Boundary-level covariates (position, word length, word log-frequency, pitch reset) are stimulus properties. Semantic shift remained the only robust predictor of residualized suppression after simultaneous adjustment for all covariates (d = −0.38, p = .004). Narrative position, lexical frequency, word length, and a prosodic boundary-strength feature (pitch reset) did not contribute. Only overall F0 *level* was marginally associated (p = .046); the targeted boundary-prosody measure (pitch reset) was not, indicating that the marginal F0 effect reflects pitch level rather than boundary prosody. (Word frequency is a within-stimulus log-count; word length, a cleaner lexical proxy, was likewise null. Boundary covariates were aligned to the stored epoch table by optimal assignment on shift, mean |Δshift| ≈ 0.012.)

| Predictor | $\beta$ (mean) | t(60) | p | d |
| --- | --- | --- | --- | --- |
| Semantic shift | −0.0030 | −2.96 | .004 | −0.38 |
| Narrative position | +0.0015 | +1.53 | .13 | +0.20 |
| Word length | +0.0001 | +0.06 | .95 | +0.01 |
| Word log-frequency | +0.0008 | +0.63 | .53 | +0.08 |
| **Pitch reset**<br>**(prosodic**<br>**boundary**<br>**strength)** | +0.0003 | +0.29 | .77 | +0.04 |
| **Pause**<br>**duration** | +0.0005 | +0.56 | .58 | +0.07 |
| **Acoustic**<br>**envelope** | −0.0007 | −0.64 | .53 | −0.08 |
| **Mean F0**<br>**(pitch level)** | −0.0021 | −2.04 | .046 | −0.26 |

**Supplementary Table S2.** Syntactic and lexical-semantic co-modeling battery. Each feature was added as a competing parametric boundary regressor (boundary×feature) alongside boundary×semantic-shift in the controlled deconvolution (baseline shift kernel d = −0.564). “Shift retained” = co-modeled shift d as a percentage of baseline; “competitor d” = the competitor’s own 0.4–1.0 s kernel. Features derived from the Stanza Hebrew pipeline: Universal-Dependencies parse for the syntactic measures, IAHLT named-entity recognition for referent introduction (N = 61). The shift kernel is robust to the dependency-structure regressors that are the closest analog to Zioga et al. (2023; dependency length 100%, tree depth 77%) and to referent introduction, content turnover and sentence length (77–101%), but is attenuated by clause and verb count, with which semantic shift is collinear (within-participant r ≈ −0.40 to −0.46). Because co-modeled betas are unstable under this collinearity, a direct variance partition of the per-boundary neural release was used: semantic shift retained a significant *unique* contribution after partialling clause count (semipartial p = .005) and verb count (p = .028), with clause/verb count contributing uniquely in the opposite direction, i.e. the release is co-determined by, but not reducible to, propositional quantity. Entering the full seven-regressor battery simultaneously attenuates the shift kernel to 42% of baseline (d = −0.238, p = .067, not cluster-significant), and the three size regressors (sentence length, clause count, verb count) alone to 40%, the expected behaviour when strongly collinear competitors are co-modelled, and the reason the variance partition rather than the co-modelled coefficients is the appropriate test here.

| **Competing**<br>**regressor** | **Type** | **Shift $d$ (co-**<br>**modeled)** | **Shift retained** | **Competitor**<br>**kernel $d$** |
| --- | --- | --- | --- | --- |
| **Mean dependency length** | syntactic (dependency structure) | −0.562 | 100% | −0.07 (n.s.) |
| **Named-entity (referent) introduction** | lexical-semantic | −0.568 | 101% | ≈0 (n.s.) |
| **Content-word turnover** | lexical-semantic | −0.459 | 81% | +0.23 (n.s.) |
| **Clause length (word count)** | length | −0.440 | 78% | +0.23 (n.s.) |
| **Tree depth** | syntactic | −0.435 | 77% | +0.24 |
| **Clause count** | syntactic / propositional quantity | −0.296 | 52% | +0.41 |
| **Verb count** | syntactic / propositional quantity | −0.211 | 37% | +0.44 |

**Supplementary Table S3.** Robustness of the burst definition. Beta-burst detection was repeated while varying the amplitude threshold (50th–90th percentile), the minimum burst duration (50–150 ms), and the detection band (12–25 vs 13–30 Hz), and the boundary burst-rate decrease and the controlled semantic-shift kernel (0.4–1.0 s Cohen’s d, co-modelling word onset, lexical surprisal and the acoustic envelope) were re-computed at each setting (N = 61). Δ burst rate is the boundary decrease as a proportion of the mean burst rate; all shift-kernel entries were sign-flip-cluster significant (p < .05). * marks the canonical definition (70th percentile, 100 ms, 12–25 Hz). The boundary decrease was significant at every threshold. It grew proportionally at stricter thresholds, but this does not discriminate a burst from a graded-power account: with a measured detector gain exponent of ∼4, a pure amplitude change necessarily produces a proportional decrease that grows as the threshold cuts further into the tail (see Results, H3). The semantic-shift kernel was essentially invariant (d ≈ −0.5 to −0.58, peak ∼+0.9 s) across all thresholds, minimum durations, and both detection bands, so the graded effect is not an artifact of the burst definition.

| Detection setting | $\Delta$ burst rate | Shift-kernel $d$ | $t(60)$ | $p$ | Kernel peak |
| --- | --- | --- | --- | --- | --- |
| A. Amplitude threshold — boundary burst-rate decrease (100 ms, 12–25 Hz) |  |  |  |  |  |
| 50th pct | –8.5% | — | –7.61 | $2.3 \times 10^{-10}$ | — |
| 60th pct | –10.6% | — | –7.04 | $2.1 \times 10^{-9}$ | — |
| 65th pct | –11.5% | — | –6.66 | $9.6 \times 10^{-9}$ | — |
| 70th pct * | –12.6% | — | –6.43 | $2.3 \times 10^{-8}$ | — |
| 75th pct | –13.6% | — | –6.09 | $8.6 \times 10^{-8}$ | — |
| 80th pct | –14.5% | — | –5.56 | $6.7 \times 10^{-7}$ | — |
| 85th pct | –15.3% | — | –4.85 | $9.0 \times 10^{-6}$ | — |
| 90th pct | –14.8% | — | –3.65 | $5.5 \times 10^{-4}$ | — |
| B. Amplitude threshold — semantic-shift kernel (100 ms, 12–25 Hz) |  |  |  |  |  |
| 60th pct | — | –0.555 | –4.34 | $5.6 \times 10^{-5}$ | +0.92 s |
| 65th pct | — | –0.559 | –4.37 | $5.1 \times 10^{-5}$ | +0.92 s |
| 70th pct * | — | –0.564 | –4.41 | $4.4 \times 10^{-5}$ | +0.92 s |
| 75th pct | — | –0.575 | –4.49 | $3.3 \times 10^{-5}$ | +0.92 s |
| 80th pct | — | –0.552 | –4.31 | $6.1 \times 10^{-5}$ | +0.92 s |
| C. Minimum duration & detection band — semantic-shift kernel (70th pct) |  |  |  |  |  |
| 50 ms | — | –0.478 | –3.73 | $4.3 \times 10^{-4}$ | +0.89 s |
| 75 ms | — | –0.548 | –4.28 | $6.9 \times 10^{-5}$ | +0.90 s |
| 100 ms * | — | –0.564 | –4.41 | $4.4 \times 10^{-5}$ | +0.92 s |
| 125 ms | — | –0.517 | –4.03 | $1.6 \times 10^{-4}$ | +0.92 s |
| 150 ms | — | –0.503 | –3.93 | $2.2 \times 10^{-4}$ | +0.92 s |
| 13–30 Hz (100 ms) | — | –0.588 | –4.59 | $2.3 \times 10^{-5}$ | +0.89 s |

## References

1. Bain M, Huh J, Han T, Zisserman A (2023) WhisperX: time-accurate speech transcription of long-form audio. Proc Interspeech 2023:4489–4493.

2. Baldassano C, Chen J, Zadbood A, Pillow JW, Hasson U, Norman KA (2017) Discovering event structure in continuous narrative perception and memory. Neuron 95:709–721.

3. Bastiaansen M, Magyari L, Hagoort P (2010) Syntactic unification operations are reflected in oscillatory dynamics during on-line sentence comprehension. J Cogn Neurosci 22(7):1333–1347.

4. Bastos AM, Lundqvist M, Waite AS, Kopell N, Miller EK (2020) Layer and rhythm specificity for predictive routing. Proc Natl Acad Sci U S A 117(49):31459–31469.

5. Benjamini Y, Hochberg Y (1995) Controlling the false discovery rate: a practical and powerful approach to multiple testing. J R Stat Soc Series B 57:289–300.

6. Black S, Gao L, Wang P, Leahy C, Biderman S (2021) GPT-Neo: large scale autoregressive language modeling with Mesh-TensorFlow. Version 1.0. Zenodo. doi:10.5281/zenodo.5297715.

7. Caspi Y, Atia B, Goldstein A (2026) Approach motivation sharpens shared neural processing: dissociable beta and alpha synchrony. Soc Cogn Affect Neurosci 21(1), nsag058. doi:10.1093/scan/nsag058.

8. Ehinger BV, Dimigen O (2019) Unfold: an integrated toolbox for overlap correction, non-linear modeling, and regression-based EEG analysis. PeerJ 7:e7838.

9. Engel AK, Fries P (2010) Beta-band oscillations—signalling the status quo? Curr Opin Neurobiol 20(2):156–165.

10. Feingold J, Gibson DJ, DePasquale B, Graybiel AM (2015) Bursts of beta oscillation differentiate postperformance activity in the striatum and motor cortex of monkeys. Proc Natl Acad Sci U S A 112(44):13687–13692.

11. Gramfort A, Luessi M, Larson E, Engemann DA, Strohmeier D, Brodbeck C, Goj R, Jas M, Brooks T, Parkkonen L, Hämäläinen M (2013) MEG and EEG data analysis with MNE-Python. Front Neurosci 7:267.

12. Hasson U, Nir Y, Levy I, Fuhrmann G, Malach R (2004) Intersubject synchronization of cortical activity during natural vision. Science 303(5664):1634–1640.

13. Hedge C, Powell G, Sumner P (2018) The reliability paradox: why robust cognitive tasks do not produce reliable individual differences. Behav Res Methods 50:1166–1186.

14. Hyvärinen A, Oja E (2000) Independent component analysis: algorithms and applications. Neural Netw 13(4–5):411–430.

15. Jas M, Engemann DA, Bekhti Y, Raimondo F, Gramfort A (2017) Autoreject: automated artifact rejection for MEG and EEG data. NeuroImage 159:417–429.

16. Jung-Beeman M (2005) Bilateral brain processes for comprehending natural language. Trends Cogn Sci 9(11):512–518.

17. Kurby CA, Zacks JM (2008) Segmentation in the perception and memory of events. Trends Cogn Sci 12(2):72–79.

18. Kutas M, Federmeier KD (2011) Thirty years and counting: finding meaning in the N400 component of the event-related brain potential (ERP). Annu Rev Psychol 62:621–647.

19. Lewis AG, Schoffelen J-M, Schriefers H, Bastiaansen M (2016) A predictive coding perspective on beta oscillations during sentence-level language comprehension. Front Hum Neurosci 10:85.

20. Lundqvist M, Rose J, Herman P, Brincat SL, Buschman TJ, Miller EK (2016) Gamma and beta bursts underlie working memory. Neuron 90(1):152–164.

21. Lundqvist M, Miller EK, Nordmark J, Liljefors J, Herman P (2024) Beta: bursts of cognition. Trends Cogn Sci. doi:10.1016/j.tics.2024.03.010.

22. Maris E, Oostenveld R (2007) Nonparametric statistical testing of EEG- and MEG-data. J Neurosci Methods 164:177–190.

23. McCarthy G, Wood CC (1985) Scalp distributions of event-related potentials: an ambiguity associated with analysis of variance models. Electroencephalogr Clin Neurophysiol 62:203–208.

24. Oostenveld R, Fries P, Maris E, Schoffelen J-M (2011) FieldTrip: open source software for advanced analysis of MEG, EEG, and invasive electrophysiological data. Comput Intell Neurosci 2011:156869.

25. Qi P, Zhang Y, Zhang Y, Bolton J, Manning CD (2020) Stanza: a Python natural language processing toolkit for many human languages. Proc 58th Annu Meet Assoc Comput Linguist Syst Demonstr:101–108.

26. Rao RPN, Ballard DH (1999) Predictive coding in the visual cortex: a functional interpretation of some extra-classical receptive-field effects. Nat Neurosci 2(1):79–87.

27. Rayson H, Moreau Q, Gailhard S, Szul MJ, Bonaiuto JJ (2026) Beta burst waveform diversity: a window onto cortical computation. The Neuroscientist 32(1):56–71.

28. Reimers N, Gurevych I (2019) Sentence-BERT: sentence embeddings using Siamese BERT-networks. Proc 2019 Conf Empir Methods Nat Lang Process (EMNLP-IJCNLP):3982–3992.

29. Rose NS, LaRocque JJ, Riggall AC, Gosseries O, Starrett MJ, Meyering EE, Postle BR (2016) Reactivation of latent working memories with transcranial magnetic stimulation. Science 354(6316):1136–1139.

30. Rouder JN, Speckman PL, Sun D, Morey RD, Iverson G (2009) Bayesian t tests for accepting and rejecting the null hypothesis. Psychon Bull Rev 16:225–237.

31. Sedley W, Gander PE, Kumar S, Kovach CK, Oya H, Kawasaki H, Howard MA, Griffiths TD (2016) Neural signatures of perceptual inference. eLife 5:e11476.

32. Seker A, Bandel E, Bareket D, Brusilovsky I, Greenfeld R, Tsarfaty R (2022) AlephBERT: language model pre-training and evaluation from sub-word to sentence level. Proc 60th Annu Meet Assoc Comput Linguist (Volume 1: Long Papers):46–56.

33. Sherman MA, Lee S, Law R, Haegens S, Thorn CA, Hämäläinen MS, Moore CI, Jones SR (2016) Neural mechanisms of transient neocortical beta rhythms: converging evidence from humans, computational modeling, monkeys, and mice. Proc Natl Acad Sci U S A 113(33):E4885–E4894.

34. Shin H, Law R, Tsutsui S, Moore CI, Jones SR (2017) The rate of transient beta frequency events predicts behavior across tasks and species. eLife 6:e29086.

35. Smith NJ, Kutas M (2015) Regression-based estimation of ERP waveforms: I. The rERP framework. Psychophysiology 52(2):157–168.

36. Spitzer B, Haegens S (2017) Beyond the status quo: a role for beta oscillations in endogenous content (re)activation. eNeuro 4(4):ENEURO.0170-17.2017.

37. Tal I, Abeles M (2013) Cleaning MEG artifacts using external cues. J Neurosci Methods 217(1–2):31–38.

38. Tinkhauser G, Pogosyan A, Tan H, Herz DM, Kühn AA, Brown P (2017) Beta burst dynamics in Parkinson’s disease OFF and ON dopaminergic medication. Brain 140(11):2968–2981.

39. Truccolo W, Eden UT, Fellows MR, Donoghue JP, Brown EN (2005) A point process framework for relating neural spiking activity to spiking history, neural ensemble, and extrinsic covariate effects. J Neurophysiol 93:1074–1089.

40. Tzourio-Mazoyer N, Landeau B, Papathanassiou D, Crivello F, Etard O, Delcroix N, Mazoyer B, Joliot M (2002) Automated anatomical labeling of activations in SPM using a macroscopic anatomical parcellation of the MNI MRI single-subject brain. NeuroImage 15:273–289.

41. Van Veen BD, van Drongelen W, Yuchtman M, Suzuki A (1997) Localization of brain electrical activity via linearly constrained minimum variance spatial filtering. IEEE Trans Biomed Eng 44:867–880.

42. Weissbart H, Kandylaki KD, Reichenbach T (2020) Cortical tracking of surprisal during continuous speech comprehension. J Cogn Neurosci 32(1):155–166.

43. Youssofzadeh V, Conant L, Stout J, Ustine C, Humphries C, Gross WL, Shah-Basak P, Mathis J, Awe E, Allen L, DeYoe EA, Carlson C, Anderson CT, Maganti R, Hermann B, Nair VA, Prabhakaran V, Meyerand B, Binder JR, Raghavan M (2022) Late dominance of the right hemisphere during narrative comprehension. NeuroImage 264:119749.

44. Zacks JM, Speer NK, Swallow KM, Braver TS, Reynolds JR (2007) Event perception: a mind-brain perspective. Psychol Bull 133(2):273–293.

45. Zioga I, Weissbart H, Lewis AG, Haegens S, Martin AE (2023) Naturalistic spoken language comprehension is supported by alpha and beta oscillations. J Neurosci 43(20):3718–3732.

